# Phylogenetically-Preserved Multiscale Neuronal Activity: Iterative Coarse-Graining Reconciles Scale-Dependent Theories of Brain Function

**DOI:** 10.1101/2024.06.22.600219

**Authors:** Brandon R. Munn, Eli Müller, Itia Favre-Bulle, Ethan Scott, Michael Breakspear, James M. Shine

**Affiliations:** Brain and Mind Centre, School of Medical Sciences, The University of Sydney, Sydney, Australia; Complex Systems, School of Physics, The University of Sydney, Sydney, Australia; Queensland Brain Institute, The University of Queensland, St Lucia, Australia; School of Psychology, College of Engineering, Science and the Environment, School of Medicine and Public Health, College of Health and Medicine, University of Newcastle, Callaghan, New South Wales, Australia

## Abstract

Brain recordings collected at different resolutions support statistically distinct signatures of information processing, leading to scale-dependent theories of brain function. Here, we demonstrate that these disparate neural-coding signatures emerge from the same multiscale functional organisation of neuronal activity across calcium-imaging recordings collected from the whole brains of zebrafish and nematode, as well as sensory regions of the fly, mouse, and macaque brain. Network simulations show that hierarchical-modular structural connectivity facilitates multiscale functional coordination, enhancing information processing benefits such as a maximal dynamic range. Finally, we demonstrate that this cross-scale organisation supports distinct behavioural states across species by reconfiguring functional affiliation and temporal dynamics at the mesoscale. Our findings suggest that self-similar scaling of neuronal activity is a universal principle that reconciles scale-dependent theories of brain function, facilitating both efficiency and resiliency while enabling significant reconfiguration of mesoscale cellular ensembles to accommodate behavioural demands.

## Introduction

Brain function emerges from the functional activity of neurons coordinated across multiple spatial and temporal scales (e.g., 1/f spectra)^1^. However, the brain is typically studied through localised apertures – e.g., cellular-scale in electrophysiology *vs*. systems-scale in neuroimaging – due to a lack of large-scale neuronal recordings^2^ and multiscale analytical techniques capable of characterising how neuronal coordination changes across spatiotemporal resolutions^3,4^. Understanding the cross-scale principles of neuronal activity can resolve the inconsistencies inherent within theories of brain function derived from scale-dependent analyses.

One such tension involves opposing theories of neural coding that argue for either efficiency or resiliency through inversely modifying redundant (shared) information between cells^5^. A sparse and efficient (minimally redundant) code can represent many distinct signals^6,7^, whereas a resilient (largely redundant) population code can readily generalise and robustly maintain function despite significant neuronal loss^8–10^. While the brain must balance these trade-offs, the extent to which functional neuronal activity favours either regime remains unclear.

Curiously, these opposing theories of brain function both have substantial empirical support, albeit from recordings of the brain oriented at contrasting spatiotemporal scales^11,12^. At the microscale, neuronal activity is uncorrelated, suggesting coding principles of efficiency^13^. Conversely, in macroscopic recordings, coarse-grained population activity – pooled across thousands to billions of neurons^14,15^ – appears strongly spatiotemporally correlated^14,15^, which advocates for resiliency. How can the brain be organised to reconcile these perspectives?

Recent advances in calcium imaging have made it possible to simultaneously interrogate the brain at the microscale (neurons) and the macroscale (whole-brain systems). Inspired by approaches in statistical physics^16^, we analyse neuronal activity across multiple spatiotemporal scales using Iterative Coarse-Graining (ICG)^17^ to resolve the theoretical schism between efficient and resilient dynamics. We apply ICG to whole-brain recordings in zebrafish (*Danio rerio*) and nematode (*Caenorhabditis elegans*), as well as sensory cortex in fly (*Drosophila melanogaster*)^18^, mouse (*Mus musculus*)^19^, and primate (*Macaca mulatta*)^20^, across multiple spatial and temporal scales. We first demonstrate that efficient and resilient activity emerges from a phylogenetically consistent, multiscale organisation principle. Next, we simulate neuronal dynamics atop a hierarchical-modular structural network that recapitulates the observed self-similar scaling. Finally, we show that the multiscale coordination is conserved across behaviours – despite significant reconfiguration at the mesoscale – suggesting multiple, scale-dependent neural codes. Our results provide a common framework for simplifying the analysis of large-scale neuronal recordings and unifying findings across scales, phylogenetically diverse species, and behaviours.

## Results

### Cross-scale transition from efficiency to resiliency in neuronal activity

ICG begins by calculating the pairwise correlation between all neurons, *N;* then greedily pairs by the temporal similarity (Pearson correlation) of their calcium fluorescence; coarse-grains by summing activity pairwise, before iterating at increasingly coarser scales, *l*. ICG thus finds dyadically increasing, coarse-grained, functionally coordinated ‘ensembles’ of neurons of size *K* = 2^*l*−1^ (cluster size; Fig. 1A). The dyadic approach provides a consistent ensemble size for comparison between recordings, while the activity pairing reveals spatially diverse ensembles (Fig. 1B) that would be missed using geometric techniques.

**Figure 1:**
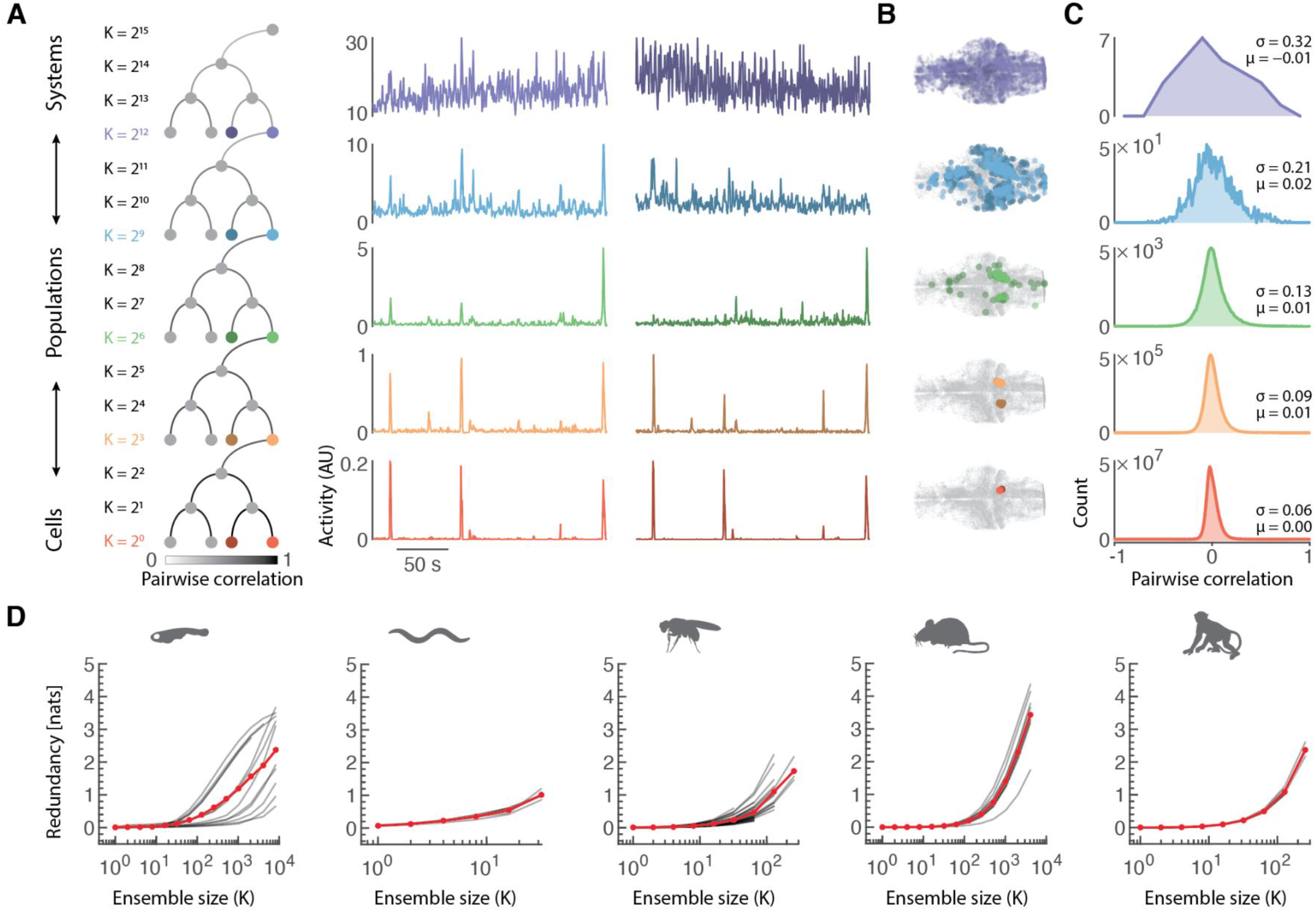
Signatures of neural coding change across scales. **(A)** Iterative Coarse-Graining (ICG) of neuronal activity is achieved by greedily pairing neurons by their pairwise correlation and summing their activity. This process is repeated on the new coarse-grained variables, iterating across larger ensembles of coordinated neurons (*K* ensemble size) and increasing the spatial scale (*l*). This approach links individual neurons to systems-level whole-brain neuronal populations across dyadically nested scales. The coarse-grained neuronal activity changes continuously from small and fast fluctuations at the neuronal scale (red) to large and slow fluctuations (purple). Pairs of combined neural ensemble activity at levels *l* = 1, 4, 7, 10, & 13 across an example branch of a zebrafish ICG hierarchy. **(B)** Examples of spatially-distributed, dyadically-increasing neuronal ensembles following ICG in the same zebrafish. **(C)** At the neuronal scale (bottom red, *l* = 1), the pairwise correlation distribution is tightly distributed around zero with few strongly correlated cells (*μ* < 10^−2^; *σ* = 0.06), whereas at the macroscale (top purple, *l* = 13), the coarse-grained neuronal ensembles correlations are broadly distributed with positive and negative correlations (*μ* = −0.01; *σ* = 0.32). **(D)** Redundancy in shared information increases with neuronal ensemble size in large-scale recordings from zebrafish (purple), nematode (blue), fruit-fly (green), mouse (yellow), and macaque (red). Faint lines are individual recordings, and solid lines are species averages. Mean (*μ*) and standard deviation (*σ*).

To demonstrate this approach, we first analyse over ∼32K neurons (∼30%) from the whole brain of larval zebrafish during spontaneous (quiescent) conditions. Analysing the ICG of neuronal activity, we can systematically track neural dynamics as they transition from microscale individual neuronal fast ‘spikes’ (Fig. 1A red) to slower macroscale fluctuations (Fig. 1A purple)^21^. Zebrafish neuronal activity shows substantial spatial coordination, with ICG revealing functional neuronal ensembles of expanding size that respect known anatomical boundaries at microscales (Fig. 1B orange) yet integrate across regions at coarser scales (Fig. 1B blue). The mean inter-cluster distance is more significant than expected if clustering by spatial proximity (p < 10^−4^), yet less than that expected of randomly paired clusters (p < 10^−4^; Fig. S1), supporting our choice to pair by temporal similarity.

ICG recapitulates known microscale-macroscale differences in pairwise correlations^13,14^. At the neuronal resolution, the correlation distribution is tightly distributed around zero (i.e., uncorrelated; example from a single zebrafish *l* = 1; mean (*μ*) = 10^−2^; standard deviation (*σ*) = 0.06; Fig. 1C, red). However, with coarsening, this distribution incrementally broadens such that at the systems level, large ensembles of spatially distributed neurons display broad correlations (*l* = 13; *μ* = −0.01; *σ* = 0.32; Fig. 1C, purple). While the ICG algorithm greedily selects positively correlated ensembles, it is notable that the approach still identifies anticorrelated populations at macroscales.

What does this cross-scale coordination imply for neural coding? To explore this, we calculated the excessive redundant shared information in pairwise neural activity estimated as the overlapping local entropy exceeding joint synergistic information (see Methods)^22^. Across all recordings and species, we observed a strong emphasis of low redundancy at the neuronal scale (*l* = 1 & *K* = 1 neurons cross-species mean ∼ 0 nats; Fig 1D). In contrast, we found a shift to increased redundancy at coarser scales (e.g., *l* = 13 & *K* = 4096 neurons cross-species mean ∼ 2.5 nats). In this way, our information-theoretic decomposition supports both efficient (neuronal) and resilient (population) neural codes. How can the same neuronal activity support these opposing functional principles?

To explain this increase in redundancy, we compare changes in the extreme values (‘tailedness’) of the pairwise correlation distribution. We calculated the kurtosis, (*μ*_4_/*σ*^4^)_*l*_, (where *μ*_4_ is the fourth central moment; Fig. 2A) across scales and found a transition from heavy-tailed correlations at the microscale to a Gaussian-like distributed (Fig. 2A green line; *μ*_4_/*σ*^4^ = 3) correlations at the macroscale. Interestingly, the scale that marked the transition to Gaussian-like kurtosis coincides with the shift from efficient to resilient coding regimes (Fig. 2B-F & Fig. 1D). Further, the decay in kurtosis across scales is slower in fish (Fig. 2B), nematode (Fig. 2C), and fruit-fly (Fig. 2D), while it is faster in mammals: mice (Fig. 2E) and primate (Fig. 2F), due to their larger cellular heavy-tailedness (*μ*_4_/*σ*^4^ > 20). We found that the few initially strongly correlated neuronal pairs drove this population-level transition – as their interdependence persisted across scales, becoming emphasised at the coarsest resolutions (Fig. S2)^4,23^. This relation supports the use of a multiscale analysis as these rare pairwise correlations are invisible through the lens of the typically reported first two moments (the mean and standard deviation). These findings suggest a multiscale coordination of neuronal activity.

**Figure 2:**
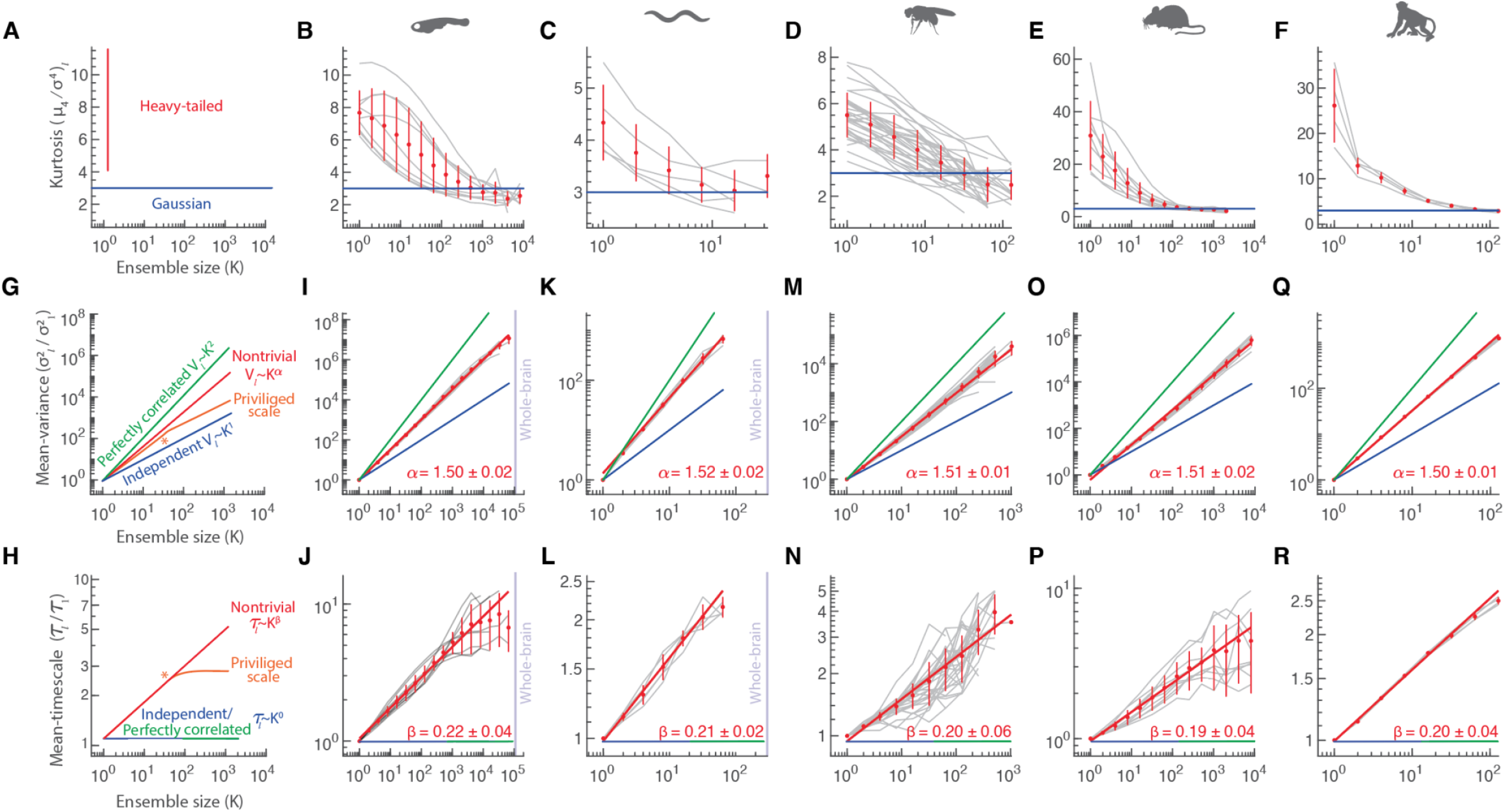
Phylogenetically-preserved self-similar scaling of neuronal activity. **(A)** The kurtosis (or ‘tailedness’) of the pairwise correlation distribution may shift across the iterative multiscale procedure from leptokurtotic ‘heavy-tailed’ (red line; *μ*_4_/*σ*^4^ > 3) to matching that of the Gaussian distribution (blue line). We calculated these measures across (**B**) zebrafish and (**C**) nematode *C. elegans* recordings with near whole-brain coverage. We also applied ICG to regional recordings from (**D**) fruit-fly *Drosophila* central brain; (**E**) mouse primary visual cortex; and (**F**) macaque primary visual and motor cortex. **(G)** The mean-variance, 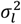, is a static measure that captures the average activity variability at each stage of the iterative multiscale procedure. The approach can differentiate between the presence of a scale-dependent transition in the neuronal organisation (privileged scales orange) and trivial scaling of either perfectly correlated (quadratic scaling - green) or independent (i.e., uncorrelated; linear scaling - blue) neural dynamics. Nontrivial self-similar scaling (1 < *α* < 2) is denoted by a power-law 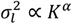 (following a straight line on logarithmic axes). **(H)** The mean-timescale, *τ*_*l*_, is a dynamic measure calculated as the average autocorrelation integral (between a log of 0 and 3 s) across scales. Trivial coarse-graining of perfectly-correlated (green) or independent (blue) signals does not alter the timescale. Nontrivial self-similar scaling (*β* > 0) is denoted by a power-law *τ*_*l*_ ∝ *K*^*β*^ (red). As above, 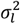 and *τ*_*l*_ were calculated across: (**I/J**) zebrafish; (**K/L**) nematode; (**M/N**) fruit-fly; (**O/P**) mouse; and (**Q/R**) macaque.

### Functional neuronal activity displays self-similar scaling preserved across species

Next, we use scaling analysis to quantify the rate of activity changes across different spatiotemporal resolutions. We compared changes in functional signatures by calculating the mean-variance (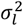; Fig. 2G) and the mean-timescale (*τ*_*l*_ estimated as the autocorrelation integral over 3 s; Fig. 2H) of the coarse-grained activity at each scale (see Methods). These measures quantify static 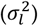 and dynamic (*τ*_*l*_) properties that can characterise the transition from sparse and fast activity at the neuronal scale to slow and variable activity at the macroscale (Fig. 1A). The rate that *σ*^2^_*l*_ and *τ*_*l*_ change across scales reflects the internal organisation of a multiscale system. We can quantify this relationship through the linear slope of *logarithmically* plotted signatures – i.e., for variance scaling as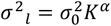, *α* can be obtained from the logarithmic log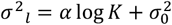.

Mean-variance and mean-timescale were chosen because their scaling is theoretically constrained, such that they can discriminate trivial coarse-graining. For instance, if the ICG is applied to an independent time series, then the mean-variance scales linearly, 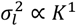, proportional to the number of independent neurons (Fig. 2G, blue), and *τ*_*l*_ ∝ *K*^0^ remains constant across coarser levels (Fig. 2H, blue; see Methods). Another trivial case arises if the neuronal activity is perfectly correlated with equivalent variability: in this case, *σ*^2^_*l*_ ∝ *K*^2^ (Fig. 2G, green) – i.e., the mean-variance scales quadratically – and again *τ*_*l*_ ∝ *K*^0^ as the timescale remains constant (Fig. 2H, green; see Methods). Alternatively, if there is a privileged scale, *K*_*p*_ (Fig. 2G/H orange-star), at which neuronal activity transitions between these states, the scaling will display a nonlinear kink. In contrast, nontrivial power-law scaling of 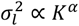 with an intermediate slope 1 < *α* < 2 (Fig. 2G red) and *τ*_*l*_ ∝ *K*^*β*^ where *β* > 0 (Fig. 2H red) indicates a nonlinear increase in variance by *α* and timescale by *β* with increasing neuronal ensemble size.

We find strong evidence supporting nontrivial, self-similar static and dynamic coordination of spontaneous neuronal activity across all species (demonstrated by a correlation coefficient of logarithmic ensemble size and variance/timescale of *r* > 0.9 across all recordings, where *r* = 1 indicates a power-law). In zebrafish, we observed power-law scaling up to 5 orders of magnitude (*α* = 1.50 ± 0.02, *μ* ± 95% CI; Fig. 2I & *β* = 0.22 ± 0.04; Fig. 2J). Similarly, we also found near whole-brain scaling in nematode (α = 1.52 ± 0.02; Fig. 2K & *β* = 0.21 ± 0.02; Fig. 2L). Furthermore, we observed power-law scaling in sensory regions of fruit-flies mushroom body (α = 1.51 ± 0.01; Fig. 2M & *β* = 0.20 ± 0.06; Fig. 2N) and within the primary visual system of mice (α = 1.51 ± 0.02; Fig. 2O & *β* = 0.19 ± 0.04; Fig. 2P) and primate (α = 1.50 ± 0.01; Fig. 2Q & *β* = 0.20 ± 0.04; Fig. 2R). The absence of a privileged scale and self-similar coherence suggests that this transition in function emerges from a systematic coordination of neuronal interdependence across coarser resolutions.

To ensure the scaling findings are not biased by the ICG algorithm and that the technique can correctly identify the trivial conditions, we repeated the analysis using generative and data-matched null models. First, we asked whether a data-matched covariance structure can explain the scaling. While this null preserved the static scaling, dynamic scaling was destroyed (Fig. S3A). Next, we tested the sensitivity of the algorithm to scale-dependencies by repeating the analysis after removing higher-order correlations outside of artificially generated scale-dependencies (e.g., *K* = 4, 32, 128), leading to scale-dependent kinks in mean-variance (Fig. S3A). We then asked whether the timescale analysis is sensitive to scaling changes by circularly time-shifting neuronal activity independently between neurons and between ICG identified neuronal ensembles (*K*_*p*_ = 2, 8, 32) before reapplying ICG, leading to trivial scaling beyond the matched scale-dependent departures (Fig. S3B).

We also asked if the intermediate static scaling emerges from a sparse sampling of a strong global coordination^24^ that can result in weak correlations at the neuronal scale. We created this generative null model by simulating neuronal activity using an inhomogeneous Poisson process with a nonstationary rate matched to an empirical coarse-grained global activity. Due to the sparse Poisson sampling, the neuronal activity appeared weakly correlated (*μ* ∼ 0.05; Fig. S4B). Nevertheless, the ICG algorithm analysis quickly revealed the global coordination (*α* = 2 & *β* = 0) within four iterations of the algorithm (Fig. S4B). Thus, neuronal activity contains a self-similarity that cannot be explained by perfectly correlated (*α* = 2; p < 1 × 10^−4^, paired t-test) or independent neuronal activity (*α* = 1; p < 1 × 10^−4^, paired t-test) and ICG scaling analysis can discriminate spurious noise-correlations, independent samplings of correlated global fluctuations, and the presence of a privileged scale-dependence.

The scaling was consistent across all animals (p > 0.05, 1-sided ANOVA with Bonferroni correction for multiple comparisons testing the null hypothesis that the scaling exponent is equal; Table S1). All scaling exponents were consistent across the first and second halves of their respective recordings (Fig. S5) and robust across alternative approaches to measure similarity, such as the Spearman coefficient and Jaccard index (Fig. S6). Therefore, these results demonstrate that a precise self-similar scaling of neuronal activity is preserved across diverse species, facilitates a breadth of functionally beneficial timescales that the system can operate across, and may reflect a whole-brain organising principle (stretching across five orders of magnitude in the zebrafish). Nevertheless, it remains to be seen where this multiscale organisation emerges from and whether this permits any information-processing benefits that may explain its preservation across broadly evolutionarily separated species.

### Multiscale structure and function and their efficiencies

How does functional self-similar scaling relate to the underlying structural connections between neurons? To address this, we analysed the ICG neuronal pairings of the *C. elegans* recordings, as both recorded cells have been uniquely identified, and the nematode’s structural connectivity (chemical synapse and gap junction) is fully characterised ^25^. Across all scales, the mean-likelihood of a functional ensemble grouping of neurons possessing a synaptic (Fig. 3A) or gap-junction (Fig. 3B) connection is more significant than chance (Fig. 3A/B, dashed lines). In particular, the grouping of ICG at finer scales correlates most strongly with structural connectivity, with 40% ± 13% of initial neuronal pairings possessing a physical connection. At coarser scales, the likelihood of all neurons possessing a structural connection is halved (∼17% ± 7%) as functional couplings become driven by higher-order (indirect) effects. These results suggest that the structural connectivity between neurons significantly informs emergent, multiscale coordinated activity.

**Figure 3:**
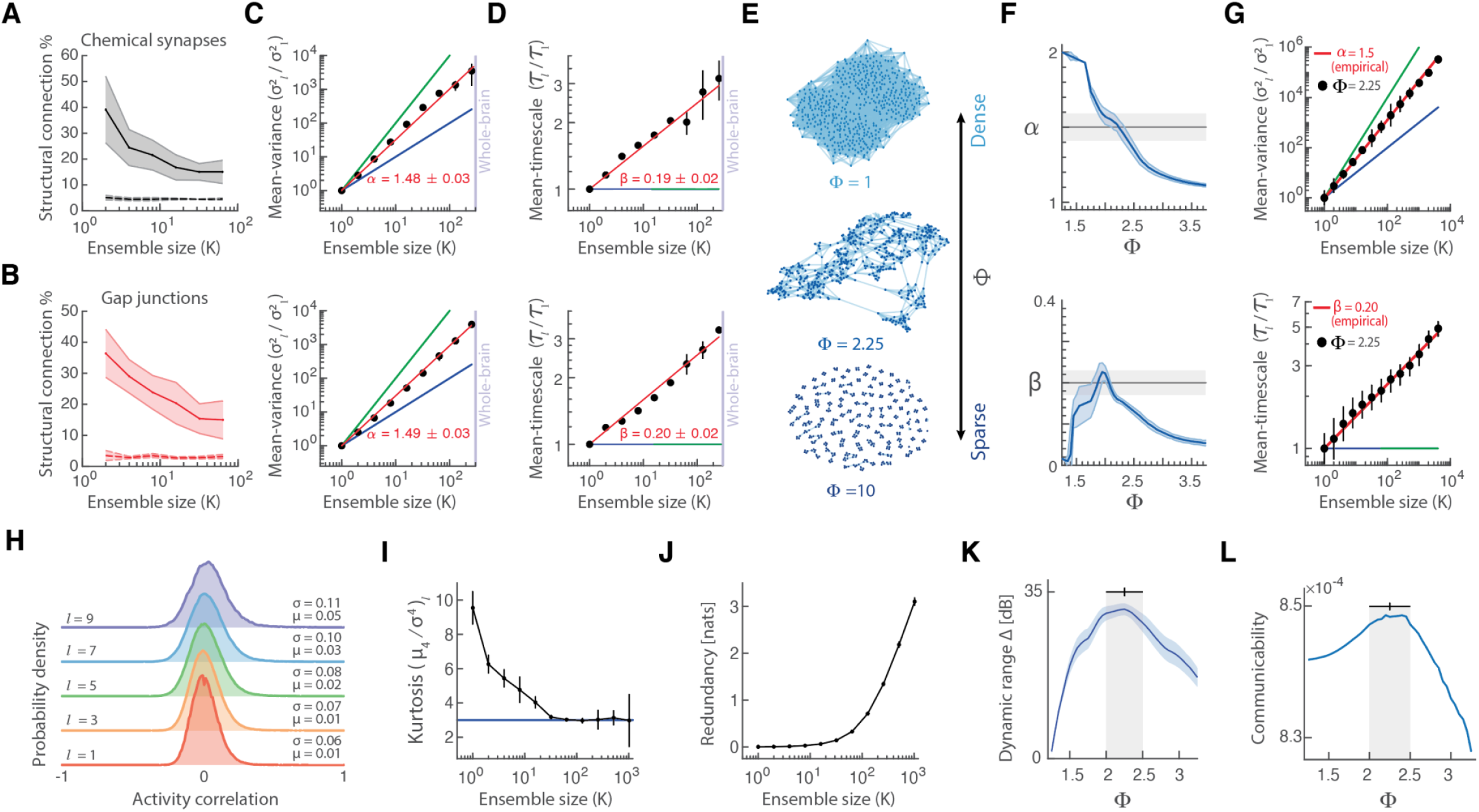
Multiscale structure and function and their efficiencies. In *C. elegans*, neurons that are structurally connected – either via chemical synapses (**A**; black) or gap junctions (**B**; red) – are likely to be functionally correlated, particularly at the microscale. Solid line: empirical recordings *μ* ± 95% CI; Dashed line: 100 random pairings. Simulations atop these physical connectivities recapitulate the empirical nematode self-similar static **(C)** and dynamic **(D)** scaling exponents (Fig. 2D). **(E)** Schematic, a hierarchical-modular network can be organised into topologies ranging from dense (Φ = 1) to sparse (Φ = 10) gradients. **(F)** Simulations on the differing self-similar topologies display proportional self-similar static and dynamic scaling that overlap with the empirical exponents at values of Φ between Φ = 2 to 2.5. **(G)** Example scaling for the fractal topology (*ϕ* = 2.25 black dots; *μ* ± *σ* over 100 simulations) with empirically observed scaling overlaid (red). **(H)** The empirical matched fractal network recapitulates the transition in pairwise correlation distribution that smoothly flows from tightly distributed around zero that broadens at coarser scales (see Fig. 1C). **(I)** The model also recapitulates the pairwise correlation distribution transition from heavy-tailed (high kurtosis) to normally distributed (kurtosis ∼ 3). **(J)** Similarly, this simulation displays increasing redundancy with neuronal population ensemble size (see Fig. 1D) **(K)** The network structures that match the empirical multiscale dynamics also possess a maximal dynamic range, Δ, the range (in dB), where variations in population activity can discernibly code variations in the input. **(L)** The network structures that match the empirical multiscale dynamics also possess peak communicability, a measure of the capacity to propagate information across the system. Error bars are *μ* ± *σ*.

To investigate the structural origin of multiscale neuronal dynamics, we next simulated cellular activity using a stochastic three-state cellular automaton neuronal model^26,27^. This branching process is a canonical model of neuronal activity^28,29^ that reduces biological complexity, improving computational efficiency (facilitating simulations across multiple topological families), and it is routinely utilised to explore universal dynamics (following that universality is independent of system details)^16,17,27,30^. Briefly, cells are simulated in discrete timesteps and can exist in either of three states: Quiescent-Active-Refractory (QAR). Cells enter the Active (‘spiking’) state from the Quiescent state either stochastically via an Active presynaptic neuron or through an external input (Poisson drive), after which the Active state cells enter the Refractory state before ultimately returning to Quiescence (see Methods). The cellular model was able to generate rich (synchronous and asynchronous) temporal patterns of coordinated neural activity and the correlation between structure-function that decays with scale (significantly exceeding chance levels, p < 0.05, 95% CI; Fig. S7). Furthermore, the model recapitulated the empirical static (Fig. 3C) and dynamic (Fig. 3D) scaling when simulated atop both the chemical synapse and gap junction structural connectivities. That is, physical connections play an integral role in generating multiscale coordination. These results confirm a multiscale structure-function relationship within the nematode and support the utility of the QAR model as a generative process to investigate the structural control of functional activity scaling relationships.

Following the relationship between structure and function within the brain, we theorised that a self-similar hierarchical-modular network could reproduce the empirically-observed functional activity scaling^31,32^. Hierarchical-modular networks possess multiple scales that are strongly connected at the base level and decrease exponentially with *P*_*l*_ = 1/Φ^*l*^, where *P*_*l*_ is the probability of a connection between ensemble pairs at scale *l* where hierarchies of modules are iteratively connected (see Methods). The fractal gradient can be transitioned between dense (Φ = 0) and sparse (Φ = 10) hierarchical networks (Fig. 3E). We found that dynamics simulated on this network yield self-similar static and dynamic scaling that smoothly changes from strongly correlated (Φ = 0 → *α* = 2) to independent (Φ = 10 → *α* = 1) activity (Fig. 3F). Within a restricted range of hierarchical-modular networks (between 2 < Φ < 2.5), the simulated dynamics recapitulate both the empirically observed static and dynamic scaling exponents (Fig. 3G). Consistent with empirical findings (Fig. 1D), the correlation distribution is tightly distributed around zero at the finest scale (*l* = 1) and broadens at coarser scales (*l* = 9; Fig. 3H). Further, this model recapitulates the empirical transition in kurtosis from heavy-tailed to normally distributed pairwise correlations (Fig. 3I) that is aligned with the transition to increasing redundancy (Fig. 3J).

Next, we simulate activity across alternative topological families to identify the relationship between structure-function and multiscale dynamics. To contrast simulations across topologies, we rescaled network connectivity strength or external input to produce a biophysically consistent maintained spike rate of ∼7Hz (see Methods). Nevertheless, we found that other topologies typically used to simulate brain network structure – such as modular, scale-free degree, geometric spatially-coupled, and small-world (Fig. S8) – could not simultaneously reproduce both the static and dynamic scaling (e.g., scale-free degree and small-world) or displayed privileged scales (e.g., modular and geometric spatially coupled). These generative models emphasise that the observed scaling is nontrivial and suggest that a limited range of hierarchical-modular topologies may explain the multiscale dynamics.

What functional benefits for information processing does this precise scaling and transition between efficiency-resiliency facilitate? This transition may reflect the capacity of the network to trade off distinct responsivity with persistent activity^33^. To explore this possibility, we drove the hierarchical-modular networks with a broad range of transient inputs of increasing frequency. We measured the population spiking output over 1s, and from this, we calculated the dynamic range, Δ (dB), defined as the 10%/90% discernible stimulus range of the input-output curve^27^. The networks reproducing empirical functional scaling coincided with a peaked dynamic range (Fig. 3K). To further verify that the network regime that matched empirical scaling facilitated increased information-processing signatures, we calculated the network communicability – an estimation of diffuse information flow^33^ – across the hierarchical-modular topologies. We found that the networks that match empirical scaling also possess peaked communicability (Fig. 3L)^34,35^. These results suggest that empirically observed functional multiscale activity can facilitate increased information flow and a susceptibility to a wide range of stimuli^36^.

### Flexible multiscale behavioural reconfigurations

The analysis thus far has focused on empirical spontaneous conditions to understand the ongoing coding regimes that constrain adaptive behaviour. To examine behaviour directly, we next investigated the functional impact of a multiscale organisation during evoked behaviours in zebrafish and mice. First, we compared ICG activity between spontaneous (data collected while fish and mice were presented with a blank screen; Fig. 4A/B/E/F, top) and evoked behaviours (visually-evoked fictive-swimming while head-fixed fish were presented with left *vs*. right phototactic stimuli and mice viewed moving visual gratings; Fig. 4A/B/E/F, bottom)^21^. Our analysis found that static variance scaling is consistently self-similar across spontaneous and behavioural states (*α*∼1.5; Fig. 4A/E). In contrast, we found that the zebrafish dynamic scaling (Fig. 4B) displays a characteristic kink (Fig. 4B, orange star) consistent with the induced timescale of the phototactic stimulation. These findings suggest that a benefit of a multiscale neuronal organisation is the capacity for neural ensembles to adapt their timescales to behavioural demands.

**Figure 4:**
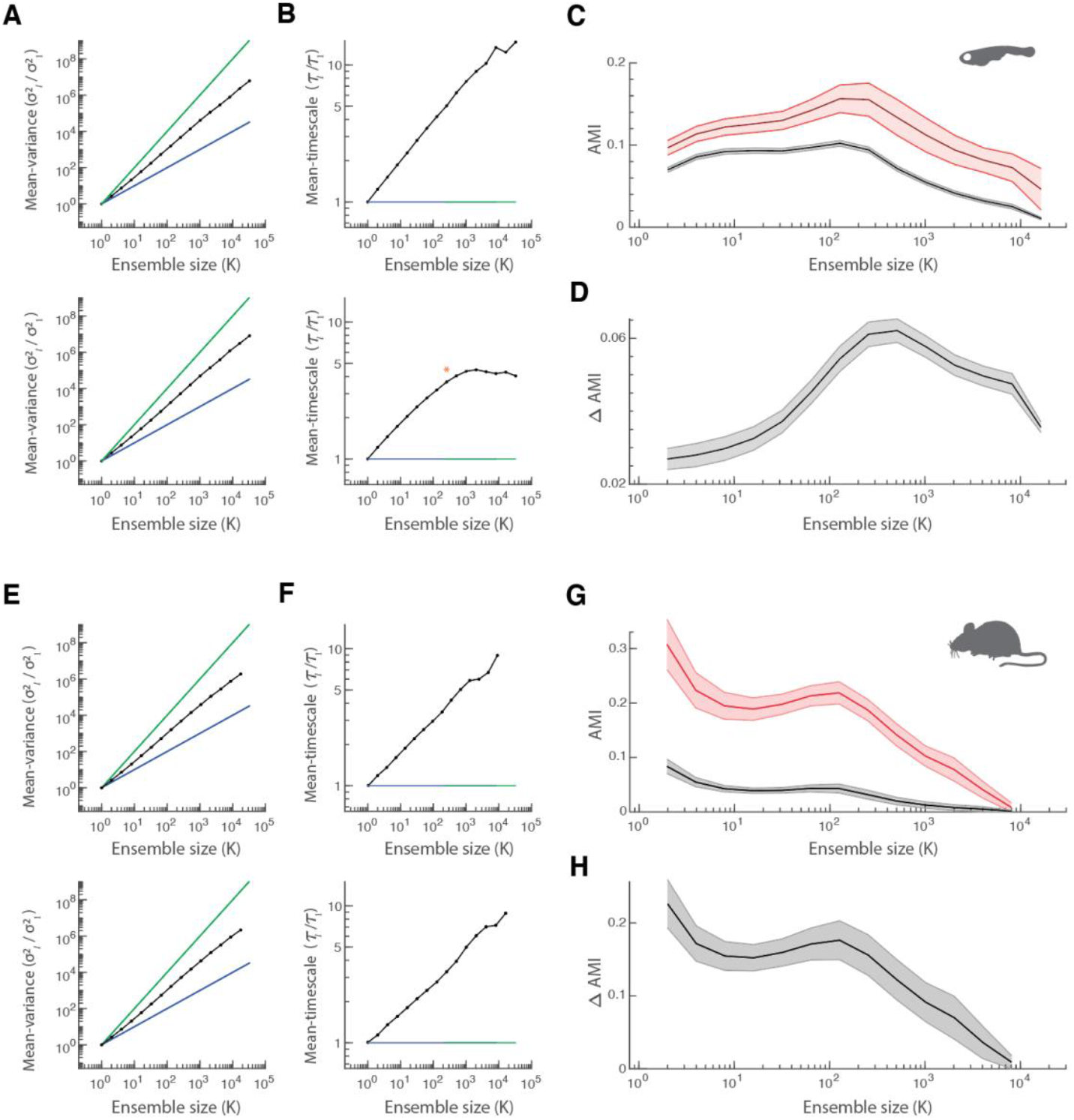
Preserved multiscale organisation during behaviour facilitates flexible scale-dependent neuronal reconfigurations across species. Comparing zebrafish multiscale neuronal activity static **(A)** and dynamic **(B)** scaling in spontaneous (top) and visually evoked phototactic alternating ‘swimming’ (bottom). Despite conserved variance scaling, the time-locked stimuli entrain a temporal dependence (orange star). **(C)** Ensembles are consistent within a state demonstrated by the adjusted mutual information (AMI) that is higher for within (red) *vs*. between (black)-behaviour comparisons for *l* = 3-14 (p < 0.001; calculated from an independent sample t-test). As the same cells were recorded across spontaneous and behavioural activity, we found that the most consistent neuronal ensembles were mesoscale neuronal ensembles. **(D)** The scale-dependent difference in AMI (ΔAMI; within minus between behaviours) shows that the strongest behavioural discriminability is at the mesoscale (*K* = 512). **(E)** Static and **(F)** dynamic scaling in mice visual cortex when viewing a blank screen (spontaneous; top) and moving sinusoidal stimuli (bottom). Mice also display a peak in mesoscale ensemble stability **(G)** and behaviour discriminability **(H)** at the mesoscale; however, there is also a significant peak at the microscale. Error bars are 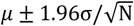.

We expected behavioural demands to mediate substantial neuronal ensemble reconfigurations. To test this hypothesis, we quantified cross-scale reconfigurations by calculating the adjusted mutual information (AMI) between neuronal ensembles (at each scale) in spontaneous and behavioural sessions (see Methods): an AMI of 1 indicates identical ensemble pairing, whereas an AMI of 0 indicates mutual information as expected from random pairings^37^. We found pairings are more stable within than between similar behaviours across split-half recordings (Fig. 4C/G. red *vs*. black;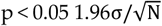). Further, we observed that functional neuronal allegiances are most stable at the mesoscale (*K*: 32 – 1024 neurons). We next calculated the difference between behaviour and rest AMI (ΔAMI), which provides an estimate of behavioural discriminability from coordinated neural activity (i.e., ensemble information content). Within the zebrafish, we found the largest ΔAMI at the mesoscale (*K* = 512; Fig. 4D). However, in conjunction with significant mesoscale discriminability, the largest ΔAMI in mouse occurred at the microscale for visual stimuli discriminability (Fig. 4H). Interestingly, this shift was discontinuous, with distinct peaks at the microscale and mesoscale. These findings support evidence that the multiscale organisation is conserved across behaviours and species, and it may facilitate simultaneous distinct neural coding principles at the micro- and mesoscale in mice.

To investigate the mesoscale networks coordinated with behaviour in more detail, we conducted a cross-scale analysis in which activity at different scales was used to predict the timing of the left *vs*. right stimuli in the zebrafish fictive swimming task^21^. We observed significant differentiation of left *vs*. right swimming behaviour across scales, with a rising proportion of significant ensembles with spatial scale (increasing from 18% at *K* = 1 to 37% explained variance at *K* = 4096) that then diminished at levels greater than *l* = 13 (i.e., peak explanatory power in ensembles of thousands of cells).

Tracking the branches of this hierarchy can provide insight into how neurons reconfigured their coordination as a function of phototaxis. At *l* = 11, four (out of 16) ensembles significantly differentiated left *vs*. right visual input, yet each involved unique anatomical regions and characteristic dynamics associated in distinct ways with the task. One ensemble (Fig. 5, blue) was dominated by hypothalamic, thalamic, and brainstem cells, displaying distinct left *vs*. right differentiation throughout the task. Another ensemble involving the telencephalon and thalamus (Fig. 5, purple) exhibited left *vs*. right differentiation later in the task. A third ensemble (Fig. 5, pink) was aligned with switches between task blocks and involved the telencephalon. The final behaviourally significant ensemble (Fig. 5, red) showed increasing responses to the task and contained predominantly brainstem neurons. Taken together, these patterns reveal the intricate and widespread neuroanatomical reconfigurations that occur in response to adaptive behaviour, suggesting potential targets for future experiments.

**Figure 5:**
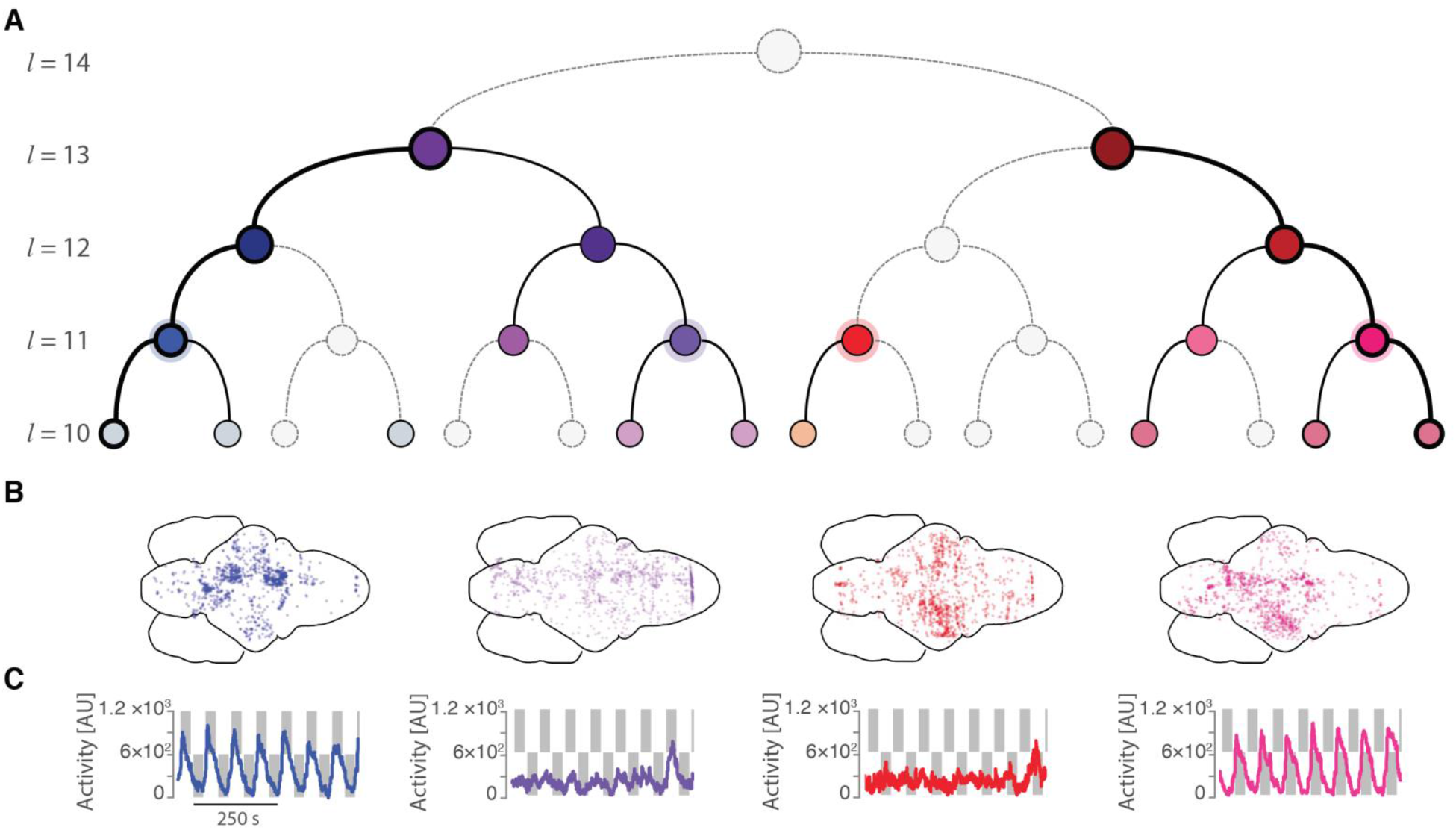
Multiscale behavioural neuronal ensemble reconfigurations. **(A)** A hierarchical tree denoting whether the activity of clusters at each level, *l*, significantly differentiated left *vs*. right phototactic stimuli: filled, coloured circles with thick lines denote significant differences between left *vs*. right at p < 0.001 (5,000 permuted null distributions), those with thin lines denote 0.001 < p < 0.05 and grey circles with dashed lines denote a lack of significant behavioural coordination (p > 0.05). **(B)** Brain-wide maps detailing the spatial coordinates of the cells that formed significant functional clusters at level *l* = 11 (outlined circles) – note the unique and spatially distributed nature of the mesoscale ensembles. **(C)** Neural activity of the significant ensembles (coloured line) with the phototactic stimulus timing (shown in grey). Note that each cluster demonstrates distinct temporal relationships with the behavioural drive.

How do these mesoscale ensembles combine to form a ‘hierarchy’ of cross-scale behavioural reconfigurations (Fig. 5)? To gain insights here, we targeted the two coarsest ensembles that maximally differentiated left *vs*. right and tracked the dyadically finer pairs of ensembles that formed these at *K* = 2048, and so on down to the cellular microscale. Using this approach, we observed several distinct tree structures, including networks that were significantly associated with the behaviour across all scales (e.g., pink and blue ensembles Fig. 5) or were transiently significant at privileged scales (e.g., purple and red ensembles Fig. 5). These results highlight how the neural assemblies responsible for distinct behaviours can fundamentally differ across scales with maximal behavioural alignment at the mesoscale.

## Discussion

We applied Iterative Coarse Graining to phylogenetically diverse and extensive calcium-imaging recordings to explore neuronal activity across multiple spatiotemporal scales. Notably, this approach revealed a self-similar organising principle that supports scale-dependent theories of brain function: at the neuronal scale, activity correlations were tightly distributed around zero with minimal redundancy, enhancing information efficiency. In contrast, when coarse-grained, the same neuronal activity exhibited broad correlations with increased redundancy, supporting resilient population coding. The transition between these two information processing regimes is marked by a shift from heavy-tailed to Gaussian-like distributed correlations. This change could be explained by a phylogenetically-conserved and nontrivial self-similar coordination of functional neuronal activity and intrinsic timescales. This functional activity scaling is closely linked to the underlying structural connectivity. Neuronal simulations on a subset of hierarchical-modular networks recapitulated the empirical scaling, which also increased the network’s information communicability and susceptibility to a broad input range. Exploration across diverse behaviours suggests that multiscale neuronal coordination may facilitate simultaneous neural coding schemas at distinct scales.

These analyses suggest that functional neuronal activity scaling is a preserved principle of neuronal organisation consistent within vertebrates (last common ancestor ∼450 million years ago) and invertebrates (last common ancestor ∼1 billion years ago)^38^. We hypothesise that the simple, functional scaling represents an evolutionary pressure of beneficial information processing – e.g., balancing neuronal efficiency and population resiliency, the susceptibility to a broad range of internal and external timescales, and the capacity to substantially reconfigure as a function of shifting behavioural demands. Future experiments should be designed to explore regional divergences in scaling exponents for insight into coding specialisation (e.g., differential scaling in the hippocampus vs sensory regions). Further, there is evidence of conserved scaling in electrophysiological recordings (across hundreds of cells), albeit with slightly shallower scaling^39^, suggesting the need for more direct comparisons across recording modalities. Finally, the structure-functional relationship we observed may explain the persistence in allometric scaling in brain-body size across species^40^.

Understanding how these cross-scale principles bridge cellular-level details and whole-brain dynamics is an important open question for neuroscience. The consistency of an invariant multiscale structure across species and simulations (even in our simplified model) provides new footholds for further theoretical modelling. For example, quantifying how the moments of neuronal correlations and activity (i.e., their mean, standard deviation, and kurtosis) change across scales provides new summary measures of system behaviour to triangulate and invert multiscale neural models. The relaxation of neural statistics from sparse and heavy-tailed correlations (leptokurtotic) at fine scales that accumulate to redundant and variable at coarse scales is a central feature of neuronal activity in all species we analyse. This shift in activity regimes motivates the need for new population models that violate the ‘diffusion approximation’ upon which classic mean-field theoretical models depend^4^. Notably, identifying fixed points under iterative coarse-graining can elucidate critical mean-field phenomena^17^. However, the observed macroscale activity patterns are inconsistent with classic avalanches – or crackling-noise – where dynamics remain bursty across all scales^41^, supporting findings showing that criticality is sufficient^42^ but not necessary for the observed multiscale dynamic^43^. Likewise, our generative cellular model rests upon a close coupling of functional activity and structure (a hierarchical network) and is not the type of microscopic branching process that yields classical avalanches. These considerations align with the framework that adaptive neural processes are a weak subcritical process^44–46^, and that avalanche dynamics are more frequently associated with pathological states such as burst suppression^47^. As such, we hope that more complex biophysical modelling approaches will extend our findings to understand healthy^3^ and pathological^48^ multiscale brain states. For example, pharmacological manipulation can test the role of the neuromodulatory ascending arousal systems control of functional activity scaling exponent and its disengagement from the underlying physical structural connectivity^3,49^.

Our results highlight the benefit of utilising a cross-scale approach to analyse the rapidly increasing scope of neuronal recordings^1,50–52^. Through this approach, we provide evidence for universal self-similar scaling of neuronal activity across the whole brain of multiple species and identify a functional benefit for this organisation (offsetting neuronal-coding efficiency and population-coding resiliency^5^), thus reconciling scale-dependent theories of brain function^6,8^. We believe that future studies incorporating this methodology – especially in rich behavioural experiments with sophisticated causal manipulations – will hasten the identification of the cross-scale neural codes required to facilitate brain function.

### Limitations of the Study

The mammalian recordings analysed in this study are of substantial size for contrasting dynamics across multiple orders of cell populations, yet (unlike zebrafish and *C. elegans* recordings) it remains to be demonstrated whether this also applies across the mammalian whole-brain. Similarly, temporal scaling analysis is limited by the recording duration and Ca^2+^ sensor resolution demonstrated by the within- and cross-species variability. As Ca^2+^ sensor and imaging technology advances, so will our capacity to test these hypotheses in whole-brain sampling^2,53^ and their consistency within recordings made with alternative modalities^54^. Likewise, while the behaviours analysed in this study are limited to visually evoked head-fixed conditions, we expect that freely moving recordings will enrich our understanding of multiscale neural reconfigurations as a function of nuanced behaviours and behavioural information processing. Further, to isolate the role of topology constraining neural dynamics in our study, we did not utilise a biophysically complex model, which significantly reduced the number of parameters in the model and allowed testing for detail-invariant universal scaling. We anticipate that more biophysically detailed models capturing neuronal nonlinearities^55^, the balance of excitation and inhibition^13^, or the role of subcortical projections^56^ will refine our understanding of additional factors that contribute to instantiating multiscale neuronal activity and further functional benefits.

### Star Methods

#### Data and code availability

All code to reproduce analysis, simulations, and zebrafish data can be obtained at (https://www.github.com/Bmunn/ICG). All other data is open and available following the details below.

### Animals and data acquisition

#### Zebrafish

The HuC:H2B–Gcamp6f zebrafish transgenic line was used for these experiments, targeting the calcium indicator GCaMP6f to the nuclei of all neurons. Three 6-day post-fertilisation larval zebrafish (*Danio rerio*) were set in a 2% low melting point agarose (Progen Biosciences), dorsal side up, and placed into the custom-made chamber. The 3D-printed 24mm^2^ chamber had a square plastic base and a 0.2mm^2^ post in each corner. 20mm^2^ glass coverslips (ProSciTech) were fixed on each side using waterproof glue (Liquid Fusion Clear Urethane Adhesive). The glass coverslips enabled light-sheet illumination from the front and one side of the animal with minimal light distortions. Agarose-set fish were mounted onto the chamber platform with additional agarose to prevent the fish from moving throughout the experiment. Once the agarose had been set, the chamber was filled with E3 medium. Whole brain calcium fluorescence imaging was performed using a custom-built selective plane illumination microscope (SPIM) to determine the neural responses. The fish were simultaneously illuminated with two planes from the front and one side and imaged at 10μm increments in the dorsoventral axis with an exposure time of 10ms. This imaging produced a 25-slice volumetric representation of the entire brain, with a 4 Hz volumetric imaging rate. Images were captured, videos were cropped, and the transverse slices were segmented to identify ROIs corresponding to individual neurons. Each of the 25 planes was motion corrected using the N*oRMCorre* algorithm, and fluorescent traces generated by calcium transients in each ROI were extracted, demixed, and denoised using the *CaImAn* package^57^. The analysis involved three fish across spontaneous and anaesthetised states (neurons = 29802, 51319, and 33882). All Zebrafish housing, breeding, larval maintenance, and experiments were performed with approval from the University of Queensland Animal Ethics Committee. The remaining zebrafish utilised in this work were obtained from open datasets^21^. The zebrafish GCaMP6f spontaneous (n = 5) and behavioural (n = 2) recordings are described in https://doi.org/10.25378/janelia.7272617. The zebrafish were imaged with a mean imaging rate of ∼2.11 Hz.

#### C. elegans

The *C. elegans* GCaMP imaging data was initially recorded and pre-processed (n = 5; full experimental detail^58^). The dataset consists of five worm spontaneous recordings without stimulation and the neurons detected in each worm in the span across the head ganglia and motor neurons and most of the sensory neurons and interneurons along with most of the anterior ventral cord motor neurons with a mean imaging rate of ∼2.9 Hz^58^. These recordings cover the worm across various behavioural regimes, from forward/backward locomotion to rolling and turning.

#### *C. elegans* connectivity structure

The *C. elegans* multiscale connectivity analysis utilised the Cook et al., ^25^ hermaphrodite gap-junction and chemical synapse connectivity matrices. Briefly, this connectivity structure was generated using electron microscope reconstruction, which led to weighted connectivity based on the sizes of the synapses.

#### Drosophila

Two-photon imaging from fruit flies (*Drosophila melanogaster*) using GCaMP6s and spontaneous activity was analysed (n=33); recordings are described in detail in ^18^ and were obtained from (https://doi.org/10.34770/gv6w-5351).

#### Mice

The spontaneous two-photon calcium recordings in mouse visual cortex using GCaMP6s in layers 2/3 excitatory neurons spontaneous (n=9) recordings are described in detail in ^19^ and were obtained from (https://figshare.com/articles/dataset/Recordings_of_ten_thousand_neurons_in_visual_cortex_during_spontaneous_behaviors/6163622). Briefly, the recordings were performed using multiplane acquisition controlled by a resonance scanner at either 2.5 or 3 Hz and the mice were free to run on an air-floating ball. Spontaneous activity was recorded in darkness or with a grey background on surrounding monitors.

#### Macaque

Awake macaque (n = 2) two-photon calcium recordings from neurons expressing GCaMP5 in the visual cortex are described in ^20^ available from (https://github.com/leelabcnbc/sparse-coding-elife2018). Awake monkeys performed a visual fixation evoked (natural images) task and spontaneous (blank screen) recordings. The recordings were made from a metal head post implanted over the dorsal portion of V1, a region representing the lower contralateral visual field at eccentricities of 2–5 deg. As these recordings did not possess continuous temporal activity (i.e., not permitting timescale analysis), we also analysed GCaMP6f recordings from the motor cortex of awake macaques (n = 2) that were obtained from (https://datadryad.org/stash/dataset/doi:10.5061/dryad.cnp5hqc4k)^59^.

### Analyses

#### Iterative Coarse Graining

We utilised an Iterative Coarse-Graining (ICG) approach to systematically cluster pairs of neurons to obtain reduced descriptions of neural activity that respected the discrete nature of individual neurons. Specifically, we clustered neurons using the similarity of their temporal activity {*ρ*_*i*_}, calculated as the Pearson’s product-moment correlation between their activity over time,

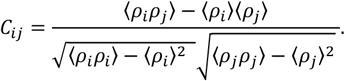

Consistent with previous approaches^17^, we greedily paired neurons by the magnitude of their correlations with removal, i.e., pairing the most positively correlated neurons followed by subsequent pairings in descending order of their correlations, such that a given neuron *i* possesses a single pairing *j*. We then coarse-grained the data by summing activity to create a new variable,

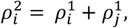

where 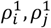 represents the neuronal activity (i.e., level 1), the new variable 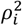 is at a lower resolution (level 2), and *i* = 1,2, … *N/*2, where *N* is the total number of neurons to be iteratively coarse-grained. This process is iterated by calculating the correlations, *C*_*ij*_, of the new variables, greedily clustering pairs, and repeating the coarse-graining,

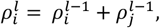

where the variable 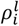 is the summed activity of *K* = 2^*l*−1^ neurons at ICG level *l*. This approach pairs neurons and thus leads to increasingly larger clusters of size *K* = 1, 2,4, …, 2^*l*−1^. We confirmed that alternative approaches to measure similarity, such as the Spearman coefficient and the Jaccard index, lead to consistent pairings.

### Statistical Measures

Statistical properties of the reduced variables at each resolution, *l*, of the system and their flow across scales are informative of the underlying mechanisms.

#### Kurtotic

We calculated the kurtosis of the pairwise correlation distribution, *C*_*ij*_, at each scale, *l*, as:

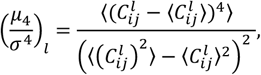

where *μ*_4_ is the fourth central moment.

#### Static

We calculated the mean-variance of the coarse-grained variables at each scale *l* by,

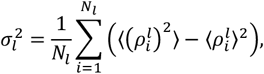

where we calculate the mean-variance across *N*_*l*_ clusters where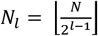. The way the variance scales across resolutions is informative of the underlying organisation of the system. If the coarse-graining procedure summed independent variables then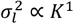, while if all variables were perfectly correlated then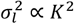. Thus, scaling following 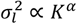with 1 < *α* < 2 is nontrivial and suggests the correlations possess an underlying self-similar coordination.

#### Dynamic

We calculated the mean-timescale of the multiscale neural activities, quantifying the temporal similarity of a signal via the mean-autocorrelation curve^60^. We first calculated the mean-normalised autocorrelation, *A*_*l*_(*t*′) as

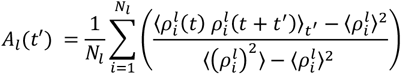

for zero and positive lags, *t*′ and at each scale *l*. We then calculated the timescale as the area under the curve (integral) estimated using the trapezoidal method 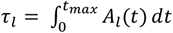 (making *τ*_*l*_ in units of time [s]), and we present results for a maximum lag of 3 seconds. We verified that the results are robust for a maximum lag between 0.5 s to 30 *s*. We then tested for scaling, *τ*_*l*_ ∝ *K*^β^. Coarse-graining two perfectly correlated signals with the same timescale or two uncorrelated signals will lead to scaling with β = 0. Hence, an exponent of β > 0 suggests nontrivial temporal scaling.

### Estimation of scaling exponent

To assess for self-similarity in the statistical properties, we fit a power-law to the static and dynamic measures across levels of *l*. Specifically, the power-law was fit from the second to the penultimate level (i.e., across *l* = 2 to *l*_*max*_ − 1). The slope fit and uncertainty were computed by linear regression in log-log space from the logarithm of the statistical measures and the logarithm of *K* across the points *K* = 2, 4, …, 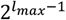 which are evenly spaced in log-log coordinates. The quality of fit to a power-law was assessed by the correlation coefficient (*r*) between the logarithm of the statistical measures and the logarithm of *K*, where *r* = 1 indicates a power-law and an r < 0.8 indicates a departure from scale-free scaling such as a privileged scale if accompanied with a characteristic kink. Note that for the most extensive dataset used in this work, the power-law fits span ∼ 5 decades, from single neuron to whole-brain zebrafish (10^5^ = 100,000 neurons over *k* = 16 dyadic scales).

### Redundancy estimation

We leveraged insights from information theory to calculate the redundant information between neuronal activity. In particular, a signal’s entropy (such as neuronal or ensemble activity) can be derived from first principles as a pointwise ‘local’ quantity that measures the information content of individual neuronal ensemble activations rather than entire sequences of activity^61^. The pointwise local entropy of a signal *X* with distinct states *x* is *h*_*x*_ = − log *P*(*x*), which quantifies the information content of a single event *x*, given the states probability *P*(*x*). Taking the expectation of this measure over all events recovers the global entropy, *H*_*X*_ = ⟨*h*_*x*_⟩ = ∑_*x*∈*X*_ *P*(*x*)*h*_*x*_. Similarly, this local approach can be extended to two signals, *X* & *Y*, with states *x* & *y*, as *h*_*x*,*y*_ = − log *P*(*x, y*). We can then apply the formalism of a joint entropy decomposition into components that are redundant, *R*, unique, *U*, and synergistic, *S*^62^, defined as

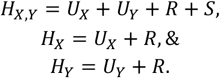

From the local measure of entropy, we can calculate an estimate of the local redundant information, 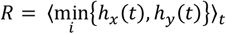, which is the minimum information that any two neurons (or ensemble of neurons) can jointly provide, averaged across the recording^61^. This estimate captures the information-theoretic basis that redundant information is the information shared between two neural signals, it is nonnegative and bound by their mutual information. In the manuscript, we present the excessive redundancy, *R*^*^ calculated as the redundant information exceeding the synergistic information, *R*^*^ = *R* − *S*, that is, the extent to which joint activity is coordinated and does not provide extra complimentary information.

### Limitations of the ICG algorithm

#### Pearson correlation

Our algorithm utilised a greedy approach based on dynamical similarity measured using the Pearson correlation of neuronal activity. As such, we are limited in the ability of the Pearson correlation to detect similarity, constrained to whether there is a linear association and lacking a causal (or directional) relationship. Furthermore, the greedy approach meant we were limited to pairwise clustering at each resolution. While both of these limitations were considered justified given that the computational efficiencies benefited from the fast calculations of pairwise correlations and greedy sorting (allowing the analysis of up to 10^5^ neurons). We also explored alternative similarity measures using the Jaccard index or Spearman correlation. We repeated our ICG analysis using these measures of similarities and found consistent scaling of our dynamics (Fig. S6).

#### Pairwise clustering: Dyads to triads

Another primary reason for pairwise clustering, instead of triadic clustering, is the nontriviality in defining a triple correlation. Nevertheless, we explored a modified triadic ICG algorithm where we sorted for the triplets of neurons with the triple correlation defined as:

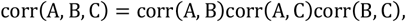

This definition is imperfect, as a single correlation near zero would significantly diminish the overall correlations, or two negative correlations would lead to an overall positive correlation. Furthermore, even this simple extension rapidly blows up the computation, taking the length of the list to be sorted from NP = (N^2^ − N)/2 where N is the number of neurons and NP is the number of unique neuronal pairings to NP choose three or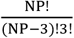. Nevertheless, utilising this algorithm on the largest sample size computationally feasible, we observe consistent static scaling between dyadic (black dots; Fig. S6) and triadic (red dots; Fig. S6) ICG after 2 iterations of triadic coarse-graining. We expect this relationship to extend to larger clustering sizes.

### Statistical null models

#### Independent vs perfectly correlated null models

To demonstrate the scaling of independent and perfectly correlated neurons, we simulated stochastic Poisson spikes with independent rates and spikes from an inhomogeneous Poisson process with an equal global rate between neurons (i.e., perfectly correlated rate but stochastic sampling).

#### Preserved scaling and artificial privileged scale data-driven null models

To ascertain whether the behaviour of these statistical properties was nontrivial and if the ICG algorithm was sensitive to signatures of a privileged scale, we generated null models from the data itself, thus preserving the covariance or temporal dynamics of all neurons. In one null model, we simulated white-noise activity with a matched covariance structure and shuffled external correlations outside paired ensembles. This null recapitulated the static scaling up to an artificial privileged scale. In another null model, we randomised the pairwise correlations by performing random circular shifts (temporal rotations) of individual neuronal activity within ensembles of neurons. These shifts rendered each cell (or ensembles of cells) independent while preserving temporal autocorrelation. This null model destroyed scaling beyond a privileged scale.

### Network construction, simulations, and analysis

#### Three-state neuronal network simulations

All networks were simulated using a stochastic spiking (cellular automata) neuronal model where a neuron, *i*, is in either one of three-state, *s*: quiescent (Q; *s*_*i*_ = 0), spiking or active (A; *s*_*i*_ = 1), or refractory (R; *s*_*i*_ = −1). The QAR stochastic model represents the basic neural model of spiking dynamics reducing biophysical complexity into three states^27,31^. This stochastic cellular model has a rich history in physics where it is used to study universal critical dynamics (as universal dynamics are independent of model details)^16,17,30^. It is computationally efficient allowing the exploration of multiple topological families (connectivity matrix *W*_*ij*_) within large-scale networks (*N* = 4096) to test the hypothesis that neuronal connectivity (structure) constrains universal multiscale activity (function).

In the model, a quiescent neuron, (Q, *s*_*i*_ = 0), will stochastically spike (A; *s*_*i*_ = 0 → 1) due to either: a stochastic external stimulus modelled as a Poisson process with transition probability P_spont_(*s*_*i*_ = 0 → 1) or evoked with probability P_evoked_(*s*_*i*_ = 0 → 1) = *W*_*ij*_ following all presynaptic cells, *j*, that spiked in the previous time step. After entering the Active state, a cell deterministically enters the Refractory period (*s*_*i*_ = 1 → −1) where it remains for two Δ*t* (so as to prevent self-reinforcing loops between mutually connected nodes) before returning to the Quiescent state (*s*_*i*_ = −1 → 0). All neurons are updated synchronously in discrete timesteps (Δ*t* = 33 ms or a sampling frequency of *f*_*s*_ = 30 Hz) and simulated for 11 seconds, with the final 10 s of spiking activity analysed with ICG (discarding initial 1 s of transience). This kind of simulation models N-methyl-D-aspartate (NMDA) spikes dynamics, which are slow currents persisting for tens of milliseconds that can nonlinearly trigger cellular spiking (acting as a superposition of many integrate and fire cells)^28^ and significantly contribute to calcium dynamics^29^. The activity was simulated on both weighted topologies and discrete binary topologies. For all simulations, we constrained the network model to have a mean firing rate of *f*_*r*_ ∼ 7 Hz by rescaling the probabilistic connectivity *W*_*ij*_ (modifying P_evoked_) so as to preserve the topological structure, however, in simulations with sparse connectivities (e.g., modular, hubless degree distributed, sparse fractal), we increased stochastic baseline drive, P_*spont*_, which is a rescaling of the external Poisson drive, *f*_spont_, as P_*spont*_ = 1 − exp(−*f*_*spont*_Δ*t*). It should be noted that other normalisation techniques may be adopted (such as preserving degree or random pruning of connections); however, we opted to preserve mean spiking dynamics with these approaches as they minimally altered the features of the topologies that we were specifically interested in exploring. We verified that pruning and randomly adding fixed strength connections with a fixed drive does not alter the qualitative findings. Nevertheless, a key limitation of this simplified model is that potentially integral biophysical constraints are not explored (e.g., the balance of excitation and inhibition as all cells are excitatory or spike-adaptation as cells possess a fixed refractory period). However, our findings demonstrate that these are not essential for the universal scaling properties observed in the simplified model and empirical data.

### Topological families

We studied five generative network models, each with a unique control parameter, allowing a broad analysis of topological properties. We considered modular networks, random rewiring networks, geometric spatially connected networks, scale-free degree distributed networks, and hierarchical-modular networks. Each distinct model parameter was simulated with 100 repetitions, and we calculated the mean and 95% confidence interval for the static and dynamic scaling to determine if there was an overlap with the experimentally observed conserved scaling. The topologies were simulated on networks of 4096 nodes, balancing sufficient iterations of ICG over 12 levels and computational limitations.

#### Modular networks

These networks were created by defining internally connected modules with a characteristic size *m*, and we present results for *m* = 2,4, …, 512. *m* = 2 implies that only pairs of neurons are connected, and the network contains 2048 modules, whereas *m* = 512 implies that the network consists of four modules. Connectivity was generated stochastically, with 95% connectivity within modules and 5% outside. Activity simulated on this network topology possessed a kink in the scaling measures at the module size (i.e., the privileged scale *K*_*p*_); we fitted the scaling exponents to scales below the module size (Fig. S8 A).

#### Random rewiring networks

These networks were created by randomly rewiring local connections within a 4096-node ring topology with mean degree = 4 and probability *λ*, which we varied in steps of 0.01 between 0 to 1. This shifts the network from an invariant ring topology (*λ* = 0) to a random topology (*λ* = 1; i.e., an Erdos-Renyi network)^32^ a rewiring probability of *λ* ∼0.2 corresponds to a small-world network (Fig. S8 B).

#### Geometric stochastic distance dependent networks

These networks were created by stochastically connecting nodes on a grid (distance of 2 between nodes adjacent nodes 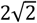 diagonally adjacent) with a distance-dependent exponential probability, *P*(*x*) ∝ *e*^−*dx*^. We presented results for *d* = 0.05 to 1.6 in intervals of 0.05. This shifted the network from local connections (*d* = 1.3; i.e., a fast drop-off in connectivity) to long-range connections (*d* = 0.05; i.e., a slow drop-off in connectivity). Interestingly, we observed a smooth transition in the initial scaling from *α* = 2 for *d* = 0.05 to *α* = 2 for *d* = 1.6, before evidence of a characteristic scale (kink in the variance) for larger clustering (Fig. S8 C; due to the characteristic scale imposed by the range of the exponential decay). Nevertheless, this initial scaling, hinted that a distant dependent exponent may be able to map between dynamical and structural scaling. Furthermore, the early match between this model and neuronal activity is promising, given the model’s important role of spatially embedded networks to generate multiscale dynamics while minimising wiring costs ^63–65^.

#### Scale-free degree-distributed networks

These networks were created by generating a scale-free degree, *d*_*eg*_, distribution with scaling exponent *γ, P*(*d*_*g*_) ∝ *d*_*g*_^−*γ*^, sampling 1,024 positive integers (0 < *d*_*g*_ *≤* 1,024), then generating the network using the Chung-Lu model^66^. We presented results for *γ* = 1.5 to 4.5 in steps of 0.1. This shifts the degree distribution from a heavy-tailed (*γ* = 1.5) to a rapidly decaying (*γ* = 4.5) short-tailed distribution (Fig. S8 D). Of note, the Barabasi-Albert model of preferential attachment^7^ possesses a scaling exponent of *γ* = 3.

#### Hierarchical-modular networks

These networks were created by defining a modular pair of connected nodes connected to other pairs with a hierarchical connection probability following *P*_*l*_ = 1/Φ^*l*^, where *P*_*l*_ is the probability of a connection between different clusters at level *l* and Φ determines the scaling^31,32^. At the beginning (*l* = 0), distinct pairs of nodes are connected, and then two groups are interconnected with a connection density *P*_*l*_ = 1/Φ^*l*^. The connection density of these nodes forms the connectivity matrix, *W*_*ij*_, referring to the probabilistic likelihood of a connection between these node groups. This process is repeated for increasing *l* and dyadically increasing group size. The value of Φ controls the hierarchical gradient. A value of Φ∼1 yields dense connectivity (i.e., all node-pairs are strongly connected across all levels), whereas Φ > 5 yields sparse connectivity as the probability of connecting node-pairs decreases exponentially across levels. We presented results for Φ = 1.25 to 3.25 in steps of 0.05. Simulations were run by converting the weighted probabilities into discrete and directed connections (randomised across iterations); nevertheless, we confirmed that the simulated results are robust when simulated using weighted connections.

#### Communicability

Communicability, *G*, is a network measure that captures how efficiently information may pass between all network nodes by pathways of all lengths (including shortest-path and higher-order connections that may be indirect between nodes)^67^. The communicability, *G*_*ij*_, between nodes *i* and *j* in a weighted network is given by:

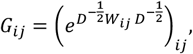

where the *m* × *m* connectivity matrix, *W*_*ij*_, is normalised by *D* = diag(*d*_*i*_) via the product 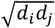 where 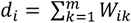 is the generalized weighted degree of node *i*. We present the summary statistic of the mean-communicability of the network, *G* = ⟨*G*_*ij*_⟩.

#### Dynamic range

To probe the information processing properties of model given stimuli, we calculated the dynamic range, Δ_s_, from the range of discernible responses to the range of stimulatory input. We calculated the response function, *F* as 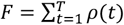, that is the spiking activity generated over *T*, where *T* = 10 s (results are robust for varying *T* = 100 ms to *T* = 2 s), in response to a transient stimulus (*T*/2 s) of strength *S*, ranging from *S* = 10^−5^ to *S* = 10^0.5^ ms^−1^, where the stimulus is modelled as afferent spikes generated as a Poisson process to each node at a stimulus rate *S*. Finally, *F* was averaged across 100 trials for each stimulus intensity. After calculating the average elicited response, F, the dynamic range, 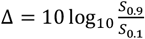, was calculated as the stimulus range (in dB) where variations in *S* can be robustly coded by variations in *F*, after discarding elicited responses that are too small to distinguish from baseline, *F*_0_, or network saturation, *F*_*max*_ ^27^. The stimulus range [*S*_0.1_, *S*_0.9_] is calculated from the elicited response range [*F*_0.1_, *F*_0.9_], where *F*_*x*_ = *F*_0_ + *x*(*F*_*max*_ − *F*_0_) which is the standard range reported ^27,68–70^.

### Adjusted Mutual Information

The adjusted mutual information quantifies the similarity between multiple partitional clustering of a set while correcting for the expected increase in mutual information for larger clusterings due to chance^37^. Given a set *S* of *N* data points, partitioned into two distinct clusterings, *U* = {*U*_1_, *U*_2_, …, *U*_*R*_} with *R* clusters, or *V* = {*V*_1_, *V*_2_, …, *V*_*C*_} with *C* clusters, where 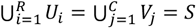 and *U*_*i*_ ∩ *U*_*j*_ = *V*_*i*_ ∩ *V*_*j*_ = ∅ for *i* ≠ *j*. The information about their overlap can be summarised from the *R* × *C* contingency table of elements 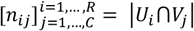, where *n*_*ij*_ denotes the number of objects that are common to clusters *U*_*i*_ and *V*_*j*_, and the partial sums of the table are 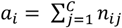 and 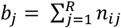. The entropy, *H*, associated with the partitioning *U* is 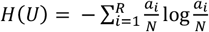 where 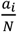 denotes the probability that a data point of *S* belongs to cluster *U*_i_ (and *v*.*v*. for *H*(*V*)) and joint entropy 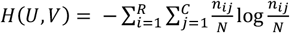. The mutual information, *MI*, between the partitions is,

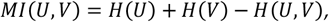

where *MI*(*U, V*) measures the reduction in uncertainty knowing a clustering tells us about the other, note that 0 < *MI*(*U, V*) *≤* max{*H*(*U*), *H*(*V*)}. The adjusted mutual information corrects the *MI* for the similarity between random clustering by removing the baseline expected mutual information, ⟨*MI*(*U, V*)⟩, under a generalised hypergeometric distribution model of randomness (i.e., stochastically), which holds that the expected value of *MI*(*U, V*) due to finite size effects and chance non-uniform joint distributions is,

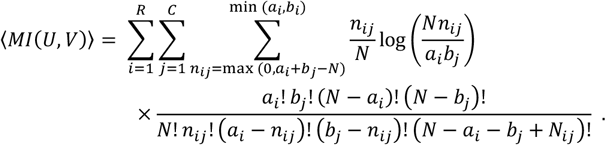

This relationship leads to the adjusted mutual information, *AMI*(*U, V*), which is calculated as,

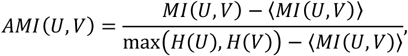

which is upper-bounded by 1, for identical partitions and equals 0, when the unadjusted *MI* equals its expected value (i.e., it is normalised in a stochastic sense).

#### General Linear Model

A mass-univariate general linear model was constructed to predict ensemble activity (at each level, *l*) during a zebrafish phototactic fictive swimming behaviour as a function of left vs. right visual signals convolved with a delayed exponential kernel (half-time = 0.4s; peak delay 0.08s) to mimic the temporal signature of the GCaMP6f indicator (as in ^71^). We generated a null distribution from the regression coefficients of 5,000 iterations of a circularly shifted dataset to estimate significance non-parametrically^72^. Significant results (p < 0.002) were plotted onto a tree denoting each level and ensemble defined using ICG on the task time series and anatomical templates warped to the Z-brain atlas^73^.

## Acknowledgements

We thank Joseph Lizier for their invaluable input concerning information-theoretic analysis.

## Authors contributions

**BRM** conceptualisation, methodology, writing, editing, investigation, analysis; **EJM** editing, methodology; **IFB** editing, investigation; **ES** editing, resources; **MB** editing, supervision; **JMS** writing, editing, supervision, funding, analysis.

**Figure S1:**
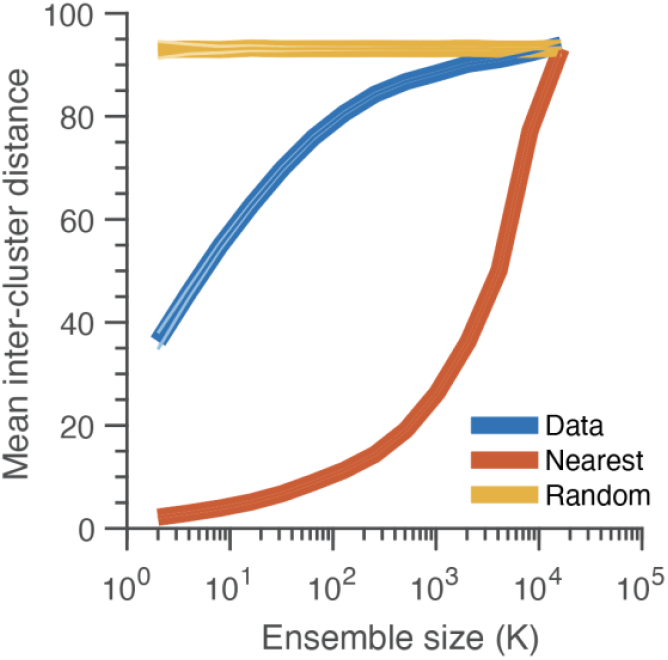
Mean inter-cluster distance for ICG ensemble size K (blue) and when clustering by spatial proximity (orange) or randomly (yellow). Error bars represent 95% CI. This result indicates that pairings display far larger spatial separations than expected for a geometric pairing.

**Figure S2:**
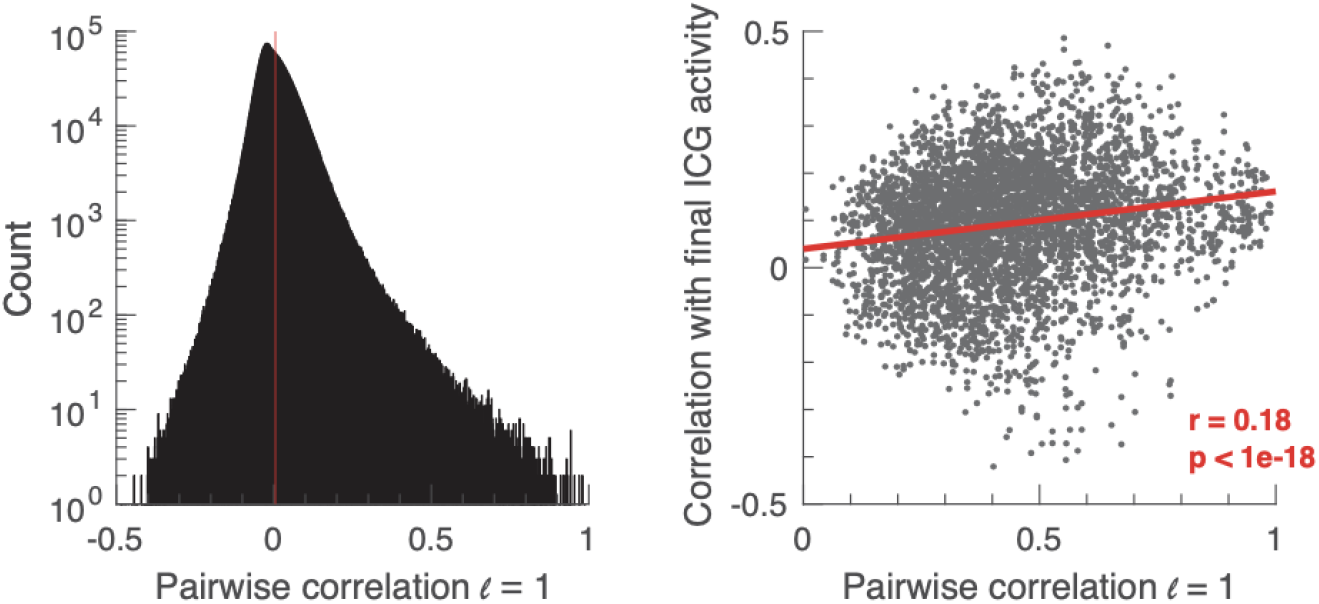
**(Left)** Pairwise correlation distribution of neurons from a zebrafish recording display a tight mean around zero with a heavy tail (note logarithmic axes). The red line indicates the smallest correlation between paired neurons following the ICG greedy algorithm. **(Right)** Individual neuronal correlations for initial ICG pairings (at *l* = 1) *vs*. the correlation between the cell and the global activity (i.e., the final ICG activity *l* = 15) suggest that the extreme initial pairings activity is preserved up to the whole brain scale.

**Figure S3:**
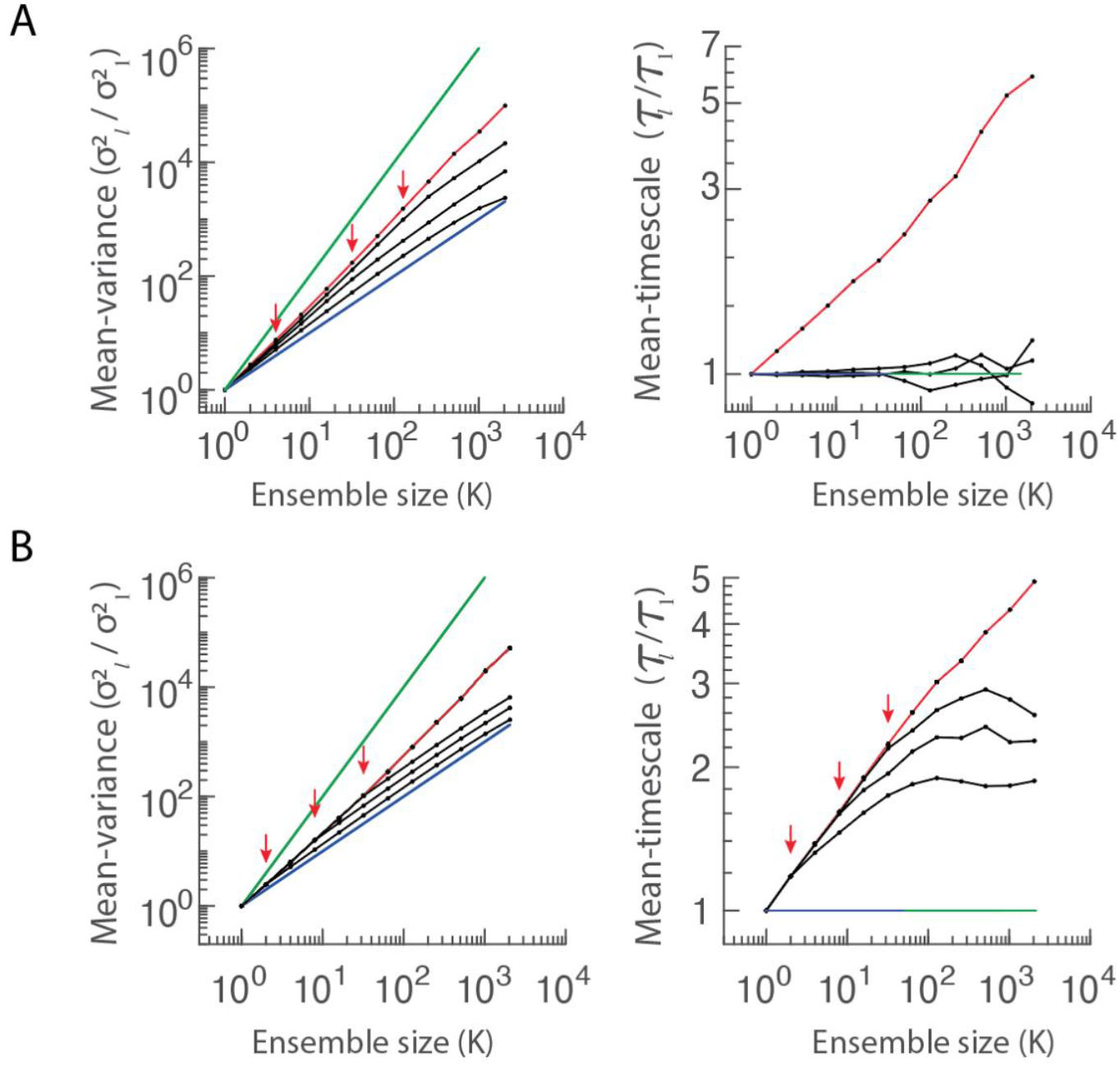
Comparison of ICG of scale-dependent static (top) and dynamic (bottom) data-matched null models. **(A)** Empirical activity (red) and artificial scale-dependent activity (curved lines examples for ICG ensemble sizes of *K* = 4, 32, 128 – red arrows). Activity is simulated atop empirical matched covariance structure after removing higher-order correlations outside of artificial scale-dependent ensemble size. (Left) Variance scales up to artificial scale dependence before scaling with a slope of unity corresponding to independent signals. (Right) The artificially generated signals do not display dynamic scaling. **(B)** We simulated null models by randomly temporally permuting (circularly shifting) empirical data within their ICG ensembles of the artificial scale-dependence – i.e., the neurons are temporally coordinated within ensembles and match empirical data yet independent outside the ensembles (examples shown for circular shifting within ensembles of size *K* = 2, 8, 32 – red arrows). This data-matched null model recapitulates the static and dynamic empirical scaling up to the artificially imposed scale dependency.

**Figure S4:**
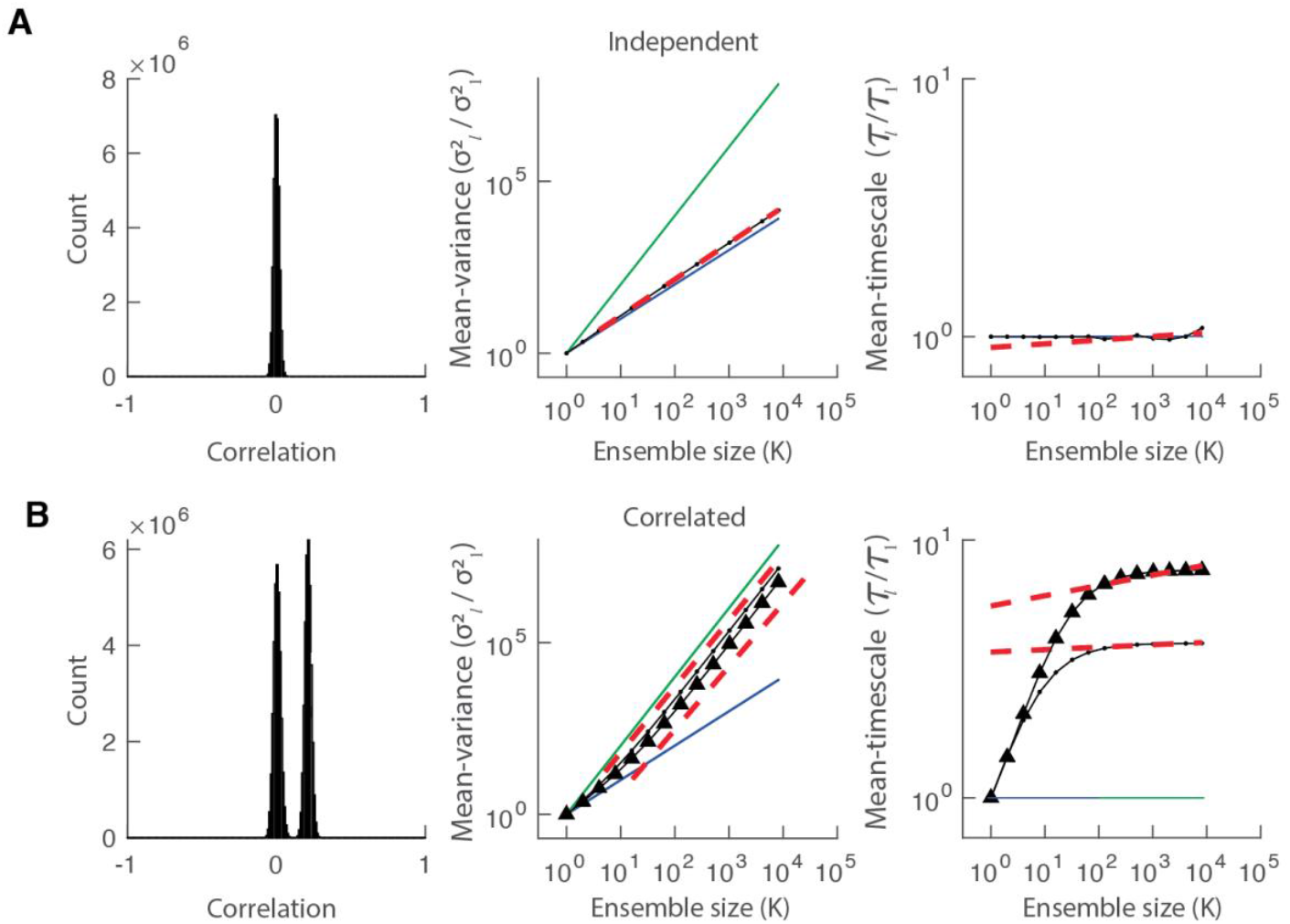
Comparison of ICG with independent and perfectly correlated Poisson spikes. **(A)** For independent Poisson-generated spikes, the pairwise correlation distribution is tightly distributed around zero (left) with static scaling exponents *α* = 1 (**c** middle) and *β* = 0 (**c** right) as theoretically expected. **(B)** Poisson spikes generated with a perfectly correlated nonstationary rate display a pairwise correlation distribution approaching zero with increasing sparse sampling of global signal (left; increasing mean pairwise correlation with mean Poisson firing rate 3.5 vs 7 Hz). Mean-variance *α* = 2 (middle) and mean-timescale *β* = 0 (right) scale as theoretically expected (3.5 Hz triangles, 7Hz circles). In the dynamic scaling, an initial transient is broader in sparse samplings.

**Figure S5:**
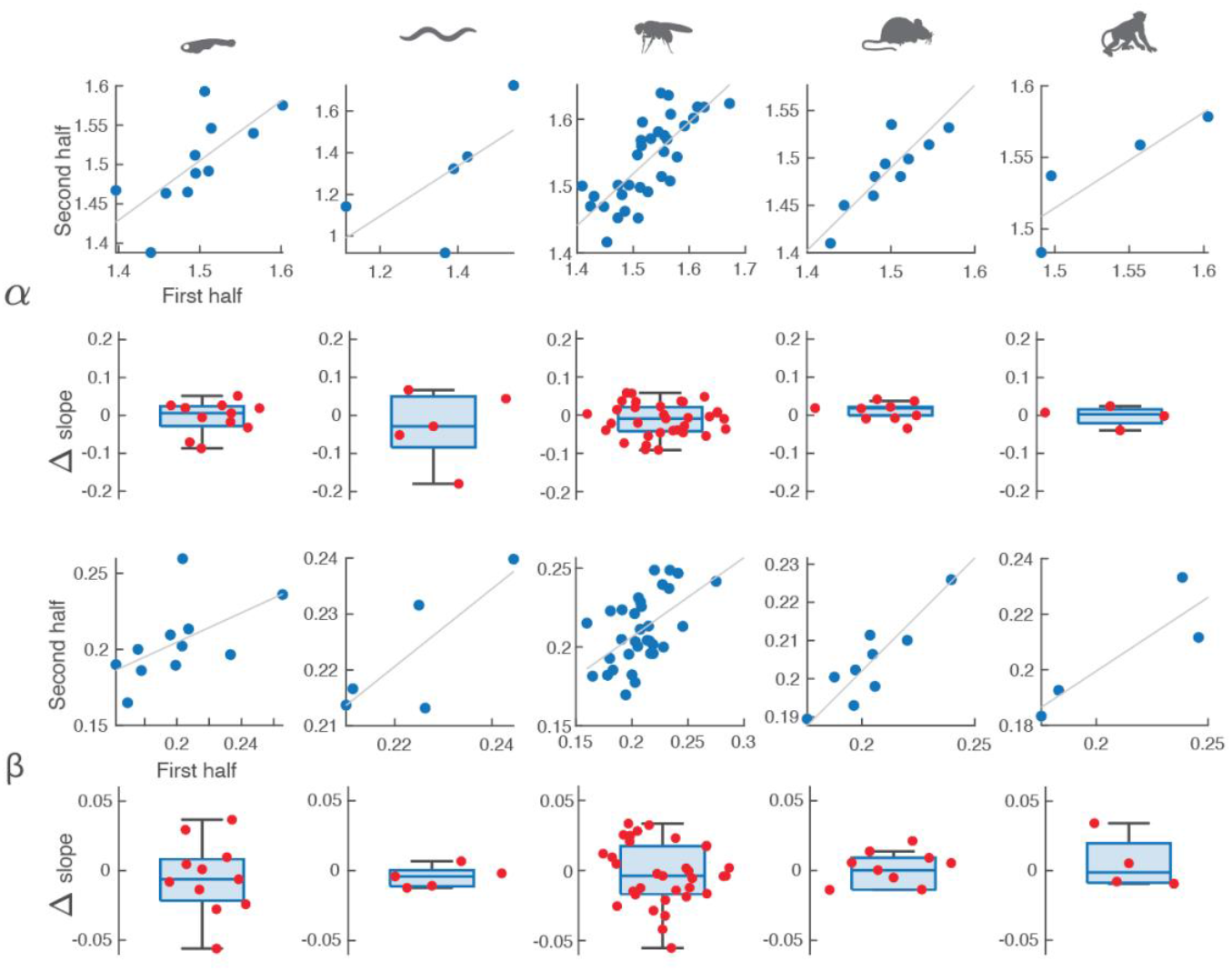
Cross-validation of scaling relationships within a recording: Mean-variance (top) and mean-timescale (bottom) scaling exponents calculated in the first and second half of each recording. The static and dynamic scaling differences (second-first) are distributed around zero, indicating an intra-difference of *α* ± 0.15 and *β* ± 0.05. Thus, the scaling is consistent within recordings of the same animal, between animals of the same species, and between species.

**Figure S6:**
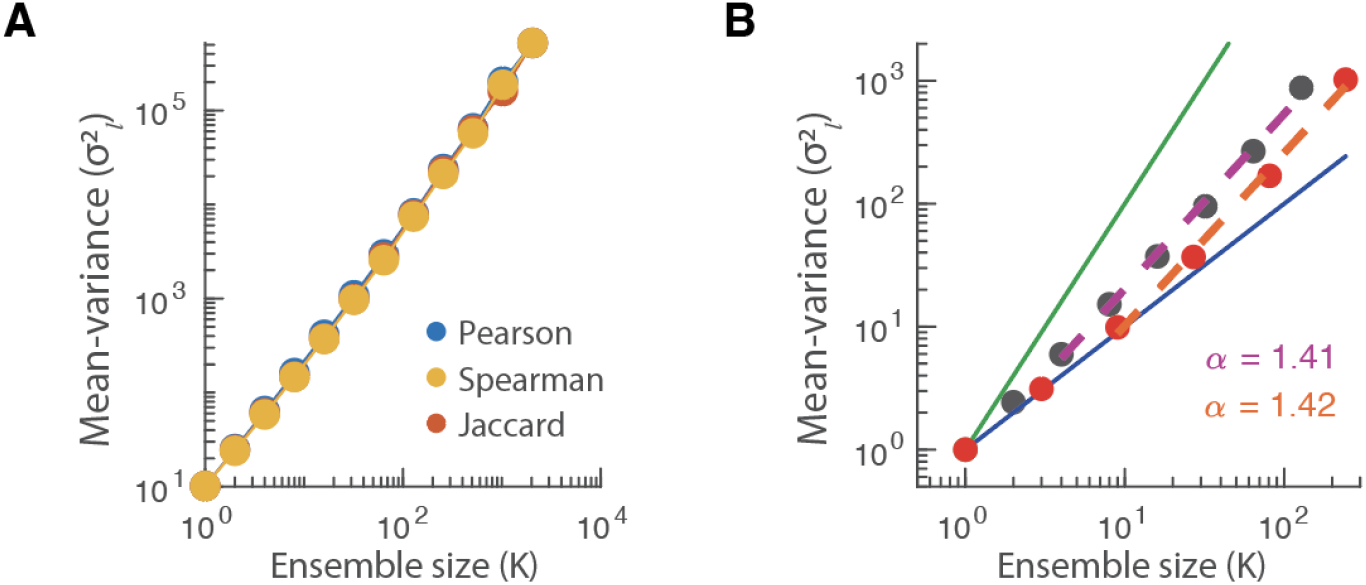
Comparison of ICG scaling using the Spearman correlation and Jaccard Index as similarity measures and triadic clustering. (**A**) The self-similar static scaling is consistent across all three similarity measures (blue-Pearson correlation; orange-Jaccard index and Pearson correlation; and yellow-Spearman). (**B**) The dyadic (black) and triadic (red) aggregating procedures produce similar scaling (dashed lines represent scale-free slope estimate purple-dyadic and orange-triadic).

**Figure S7:**
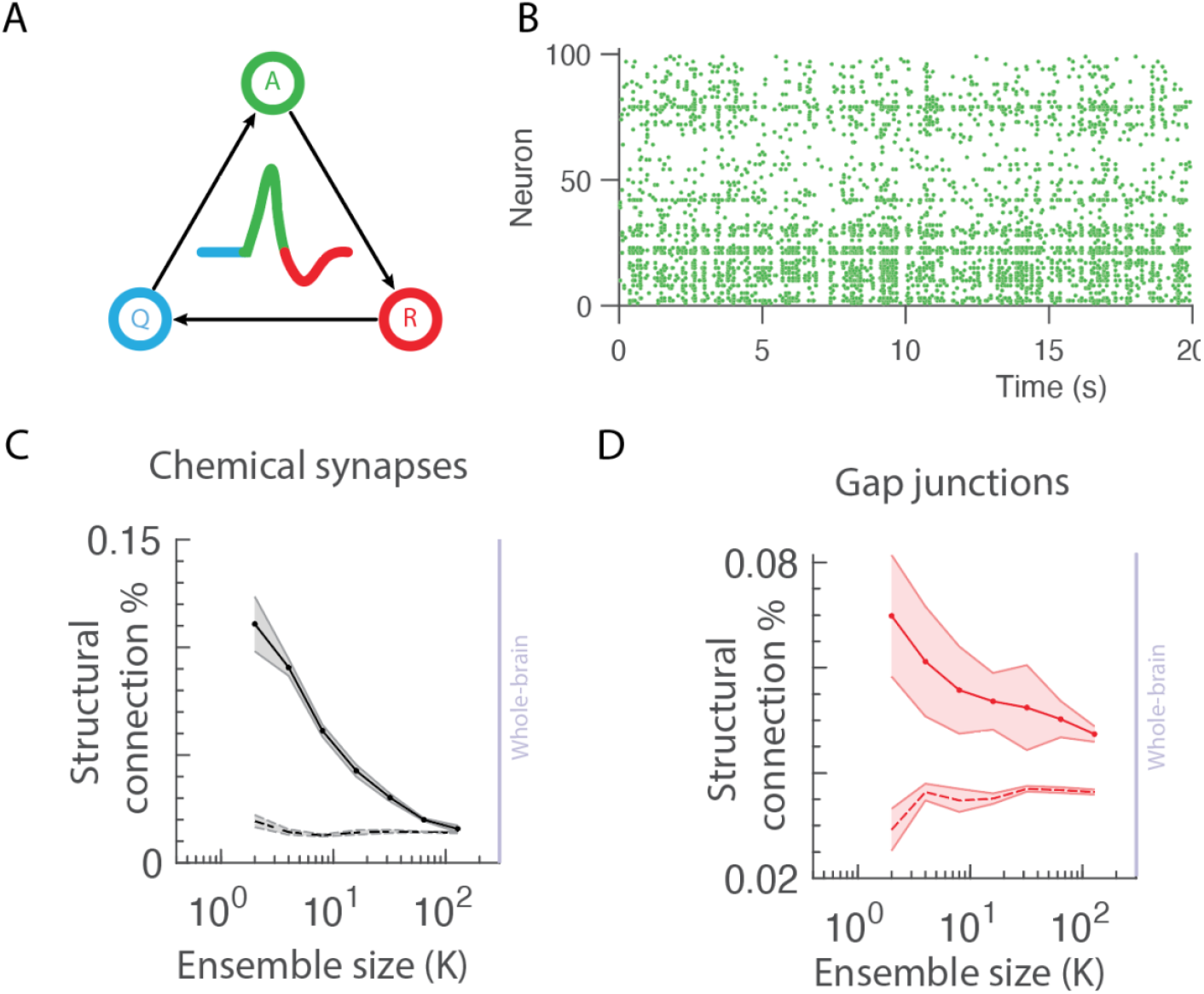
Three-state cellular model atop *C. elegans* structural connectivity. **(A)** QAR model recapitulates the three stages of an action potential (spike), Quiescence (able to spike), Active (spiking), and Refractory period (unable to spike). (**B**) Example spiking activity from the model simulated atop the *C. elegans* chemical synapse (examples of simultaneous synchronous and asynchronous activity. Likelihood of simulated activity functional connections obtained using ICG possessing a physical **(C)** chemical synaptic or **(D)** Gap junction connection.

**Figure S8:**
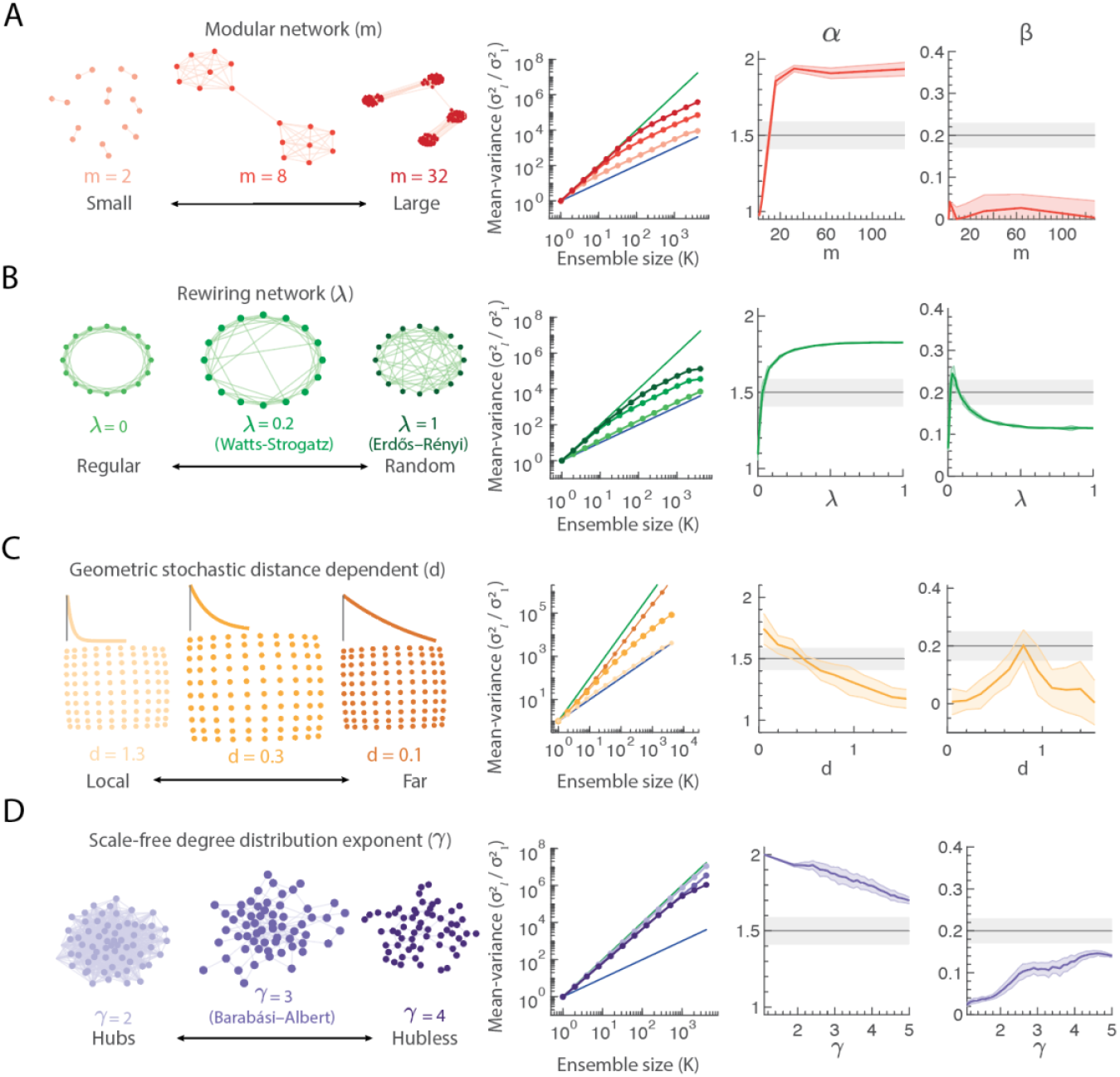
Simulated neuronal dynamics connected via different topological network families using the QAR model and their static (*α*) and dynamic (*β*) scaling exponents. **(A)** Simulations on a modular network topology do not display power-law scaling, possessing a kink above the module size. The dominant scaling exponent for module size (m; right *μ* ± *σ* across 100 simulations red shaded bar) does not coincide with the empirically observed variance exponent (black line ± grey error bars). **(B)** Rewiring topologies extending from regular (rewiring probability *λ* = 0) to random (*λ* = 1; Erdős–Rényi) via a Watts-Strogatz small-world connectivity (*λ* = 0.2), support self-similar scaling for minor rewiring’s that do not coincide with the empirically observed exponents. **(C)** Geometric stochastic distance-dependent networks where spatially embedded neurons are stochastically connected proportional to a distance-dependent exponential distribution; the slope of this decay extends the networks from locally connected (*d* = 1.3) to broadly connected to spatially far neurons (*d* = 0.1). This network recapitulates the empirical scaling from efficient to robust, albeit for a transient regime, before scaling independently beyond a finite ensemble size proportional to the range of the geometric coupling. **(D)** Scale-free degree distributed networks with degree exponent (−*γ*), including the Barabasi-Albert network (*γ* = 3), support self-similar scaling but with strong correlations even following connectivity reweighting that do not coincide with the empirically observed exponents.

**Table 1:** Table of p-values testing the hypothesis, (H_0_), that the species mean static (*α*) and dynamic (*β*) scaling exponent are equal ANOVA Tukey-Kramer corrected for multiple comparisons. The hypothesis that the scaling exponents are conserved across species is rejected at the 95% confidence interval if the p-value < 0.05.

|  |  | Static (p-value) | Dynamic (p-value) |
| --- | --- | --- | --- |
| Species A | Species B | $H_0: \langle \alpha_A \rangle = \langle \alpha_B \rangle$ | $H_0: \langle \delta_A \rangle = \langle \delta_B \rangle$ |
| Zebrafish | *C. elegans* | 0.92 | 0.92 |
| Zebrafish | Drosophila | 1 | 0.99 |
| Zebrafish | Mouse | 1 | 0.99 |
| Zebrafish | Macaque | 0.83 | 0.77 |
| *C. elegans* | Drosophila | 0.85 | 0.97 |
| *C. elegans* | Mouse | 0.88 | 0.99 |
| *C. elegans* | Drosophila | 1 | 0.99 |
| *C. elegans* | Macaque | 0.99 | 0.99 |
| Drosophila | Mouse | 1 | 0.99 |
| Drosophila | Macaque | 0.74 | 0.84 |
| Mouse | Macaque | 0.78 | 0.95 |

## Notes

### Competing Interest Statement

The authors have declared no competing interest.

